# Single-cell Transcriptomics reveals multi-step adaptations to endocrine therapy

**DOI:** 10.1101/485136

**Authors:** Sung Pil Hong, Thalia E. Chan, Ylenia Lombardo, Giacomo Corleone, Nicole Rotmensz, Giancarlo Pruneri, Kirsten R. McEwen, R. Charles Coombes, Iros Barozzi, Luca Magnani

## Abstract

Resistant tumours are thought to arise from the action of Darwinian selection on genetically heterogenous cancer cell populations. However, simple clonal selection is inadequate to describe the late relapses often characterising luminal breast cancers treated with endocrine therapy (ET), suggesting a more complex interplay between genetic and non-genetic factors. Partially, this is due to our limited understanding on the effect of ET at the single cell level. In the present study, we dissect the contributions of clonal genetic diversity and transcriptional plasticity during the early and late phases of ET at single-cell resolution. Using single-cell RNA-sequencing and imaging we disentangle the transcriptional variability of plastic cells and define a rare sub-population of pre-adapted (PA) cells which undergoes further transcriptomic reprogramming and copy number changes to acquire full resistance. PA cells show reduced oestrogen receptor α activity but increased features of quiescence and migration. We find evidence for sub-clonal expression of this PA signature in primary tumours and for dominant expression in clustered circulating tumour cells. We propose a multi-step model for ET resistance development and advocate the use of stage-specific biomarkers.

## Introduction

The outgrowth of primary luminal breast cancer (BCa) is driven by non-mutated estrogen receptor α (ERα), with all patients receiving adjuvant endocrine therapy after curative surgery (ET). This strategy significantly delays clinical relapse but does not abrogate it completely, as about ∼3% of the patients each year come back with overt relapse, inevitably leading to further metastatic development^1-3^. The frequency of relapse remains constant up to 20 years after surgery making ET-resistance the most critical clinical problem for the management of these patients^4^. The processes of adaptation and selection leading to late relapse are currently poorly understood and should be interpreted in light of adjuvant therapies.

Recent developments in next-generation sequencing (NGS) revealed that tumours are genetically heterogeneous^5-7^ and in some cancer types, heterogeneity correlates with the likelihood of recurrence and development of drug-resistance^8,9^. In some instances, targeted therapy can lead to the rapid expansion of genetically defined pre-existent resistant cells that can be explained by simple models of clonal selection^10-12^. However, this same model is mostly inconsistent with the decade-long latency observed in luminal BCa. In addition, despite recent studies showed that the majority of the genetic lesions in BCa are accumulated during the early phases of tumour development^5,13^, they failed to identify any major driver associated to metastasis and resistance, with the exception of a minor fraction of cases showing either *ESR1* mutations or *CYP19A1* amplification^14-17^. Yet, the transcriptomes of the resistant cells are profoundly heterogeneous and different from those of the primary tumour^18-20^, suggesting a contribution of non-genetic mechanisms^21^.

Rare phenotypic subpopulations, showing features of drug-tolerance and sometimes of quiescence, have been found in primary melanomas^22^, leukaemia^23^, non-small cell lung cancer^24^ and triple negative breast cancer (TNBC)^25^. In primary melanoma, a rare, transient subpopulation expressing resistant markers at high levels can survive and persist to become stably resistant^26^. Nevertheless, it remains unclear how genetic and non-genetic components contribute to different types or stages of ERα-BCa.

In this study, we used a combination of live cell imaging, single cell RNA-sequencing (scRNA-seq) and machine learning to dissect the phenotypic heterogeneity and plasticity of ERα-positive BCa and leveraged this information to identify a subpopulation of rare, pre-adapted cells both *in vitro* and *in vivo*. These cells (termed PA, from Pre-Adapted) display a unique transcriptional signature with features of dormancy and mixed epithelial and mesenchymal traits, and which was found dominant in clustered circulating tumour cells. PA cells showed a significant survival advantage under short-term ET but required further transcriptional reprogramming and genetic alterations to acquire full resistance and re-establish a proliferative phenotype *in vitro*. These results highlight the multi-faceted effects of ET at single cell level and suggest a multi-step mechanism of drug resistance that involve both non-genetic and genetic contributions.

### Phenotypical equivalent of fully resistant clones is absent in treatment-naïve cells

In order to study the dynamic process of ET resistance, we exploited an *in vitro* system that maximises reproducibility while minimising confounding factors^15,27^. Long-term oestrogen deprived (LTED) cells originate from ESR1 wild-type MCF7 that have been deprived from oestradiol for one year. This model is generally considered a good proxy to study the effect of aromatase inhibitors (AI) (Fig. 1a). We previously showed that resistance in this model is driven by amplification of the aromatase gene (CYP19A1) in combination with endogenous cholesterol biosynthesis, but not by mutated ESR1^15,28^. Even in this accelerated model, fully resistant cells emerge between 6 to 12 months of oestrogen deprivation^29^, which is incompatible with clonal selection of a pre-resistant cell^30^. In line with this, less than 1.5% of early-stage BCa show evidence of a pre-existent ESR1-mutant clone^31^ (Supplementary Table 1), suggesting key driver mutations to be acquired at a later stage. However, this model does not fully exclude pre-existence of transcriptomic clones with features of resistance. To investigate this, we generated scRNA-seq high-quality profiles for >1,200 MCF7 and >1,900 LTED cells (Supplementary Table 2).

**Figure 1.**
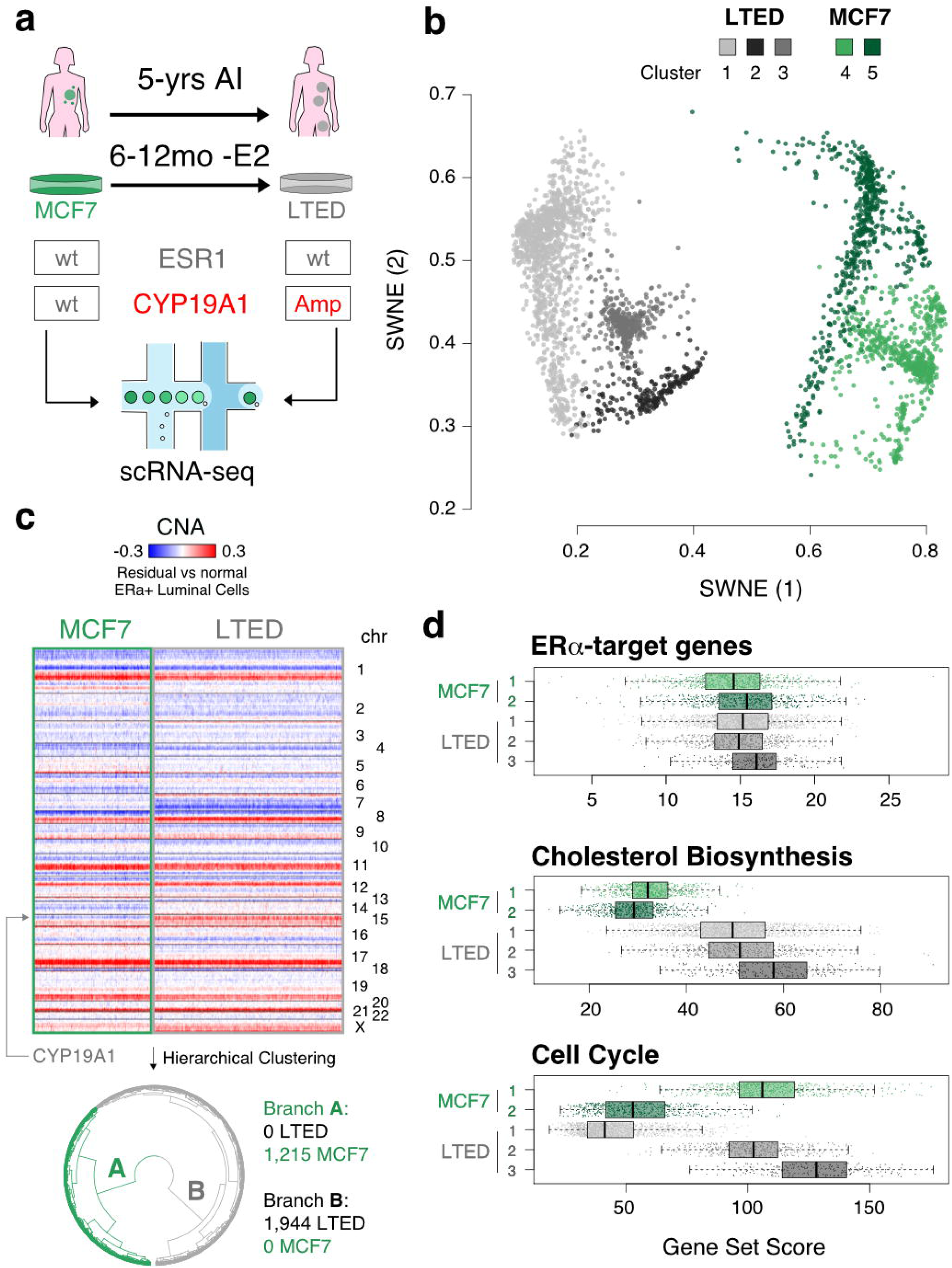
Phenotypical equivalent of fully resistant clones is absent in treatment-naïve cells. **(a)** Schematic representation of the *in vitro* approach (bottom), which mimics the development of resistance to aromatase inhibitors (AI) in patients. **(b)** Bi-dimensional representation of 3,159 single cell transcriptomes (1,125 MCF7 and 1,944 LTED) (SWNE; *k* = 16). **(c)** Copy number profiles of the cells shown in (b), as estimated from scRNA-seq profiles. Data shown as heat map and as dendrogram (hierarchical clustering; Ward’s method; Euclidean distance). **(d)** Overall expression for selected gene sets, by cluster of cells (as defined in b).

Dimensionality reduction (Similarity Weighted Nonnegative Embedding, or SWNE)^32^ showed MCF7 and LTED as completely separated populations, with no single MCF7 clustering with LTED cells (Fig. 1b). Studies in melanoma and TNBC suggest that drug-resistant cells can rapidly emerge^25,26^. This implies that in drug-naïve tumors, at least a few cells have a transcriptional profile similar to that of fully resistant cells. However, our data suggest this is not the case in luminal breast cancer cell lines, which is concordant with the long latency taken by ET-resistance to occur in most patients treated with endocrine therapies. To completely exclude any contribution of a pre-existent genetic clone, we inferred single-cell, copy number alterations (CNAs) from scRNA-seq data (Methods). Clustering of single MCF7 and LTED cells based on the inferred patterns of CNAs identified two clades, one including all the MCF7 and one all the LTED cells (Fig. 1c). In line with CYP19A1 significantly contributing to AI resistance *in vivo* and *in vitro*^15^, an amplification involving the region was found in almost all LTED cells but not in MCF7 (Fig. 1c). This was confirmed by shallow whole genome sequencing (Supplementary Fig. 1a).

Taken together, these data support that AI resistance is not driven by a pre-resistant clone (whether genetic or in a particular transcriptional state), suggesting a multi-step adaptation process in which the necessary hits occur with a different timing during ET.

Using scRNA-seq, we were able to confirm that certain pathways, such as cholesterol biosynthesis, are profoundly reprogrammed by ET at the population level (Fig. 1d and Supplementary Fig. 1b). Nevertheless, clustering of single-cell profiles identified five distinct groups (two for the MCF7 and three for the LTED), reflecting a certain degree of diversity for the activity of these pathways. This led us to further dissect the phenotypic heterogeneity of breast cancer cells with the aim of pinpointing rare, transcriptionally-defined clones in the drug-naïve condition.

### Characterisation of a phenotypically distinct population of plastic cells in luminal breast cancer

We first aimed at identifying any surface marker that could guide the dissection of the phenotypic heterogeneity of luminal breast cancer cells. Genes showing high transcriptional variability across single MCF7 cells were intersected with annotation in the Cell Surface Protein Atlas^33^. Among the top 30 genes, we identified CD44 (Supplementary Fig. 2) which has been previously recognised as a marker of plastic cells in various solid tumours^34-36^. Further investigation confirmed variable expression of CD44 in primary tumours (Supplementary Fig. 3a-c and Fig. 2a,b). CD44 positive cells were found significantly enriched after treatment in a cohort of neo-adjuvant AI patients (Fig. 2a; 3.8-fold, *p*-value = 0.0032; two-tailed paired *t*-test) as well as in matched AI-treated primary-metastatic (Fig. 2b and Supplementary Fig. 3d; 2-fold, *p*-value = 0.0029; Wilcoxon signed-rank test), suggesting higher chances of survival to ET for CD44-expressing cells *in vivo*. We next sought to investigate if CD44^high^ cells can be also found at other active sites in breast cancer patients. Interestingly, we found substantial CD44^high^ cells in pleural effusions from all four patients examined (Supplementary Fig. 3e). In line with this, the fraction of FACS-sorted CD44^high^ cells was significantly increased in LTED (upper panels in Supplementary Fig. 3f,g). Extensive functional characterisation of these cells demonstrated that MCF7-CD44^high^ cells were more invasive, more clonogenic, and could form first and second-generation of mammosphere at higher efficiency than CD44^low^ cells (Supplementary Fig. 3h-j). In agreement with previous studies^36^, CD44^high^ cells also showed cellular plasticity as they could recapitulate the entire population while CD44^low^ were capable of generating only CD44^low^ cells (Supplementary Fig. 3f).

**Figure 2.**
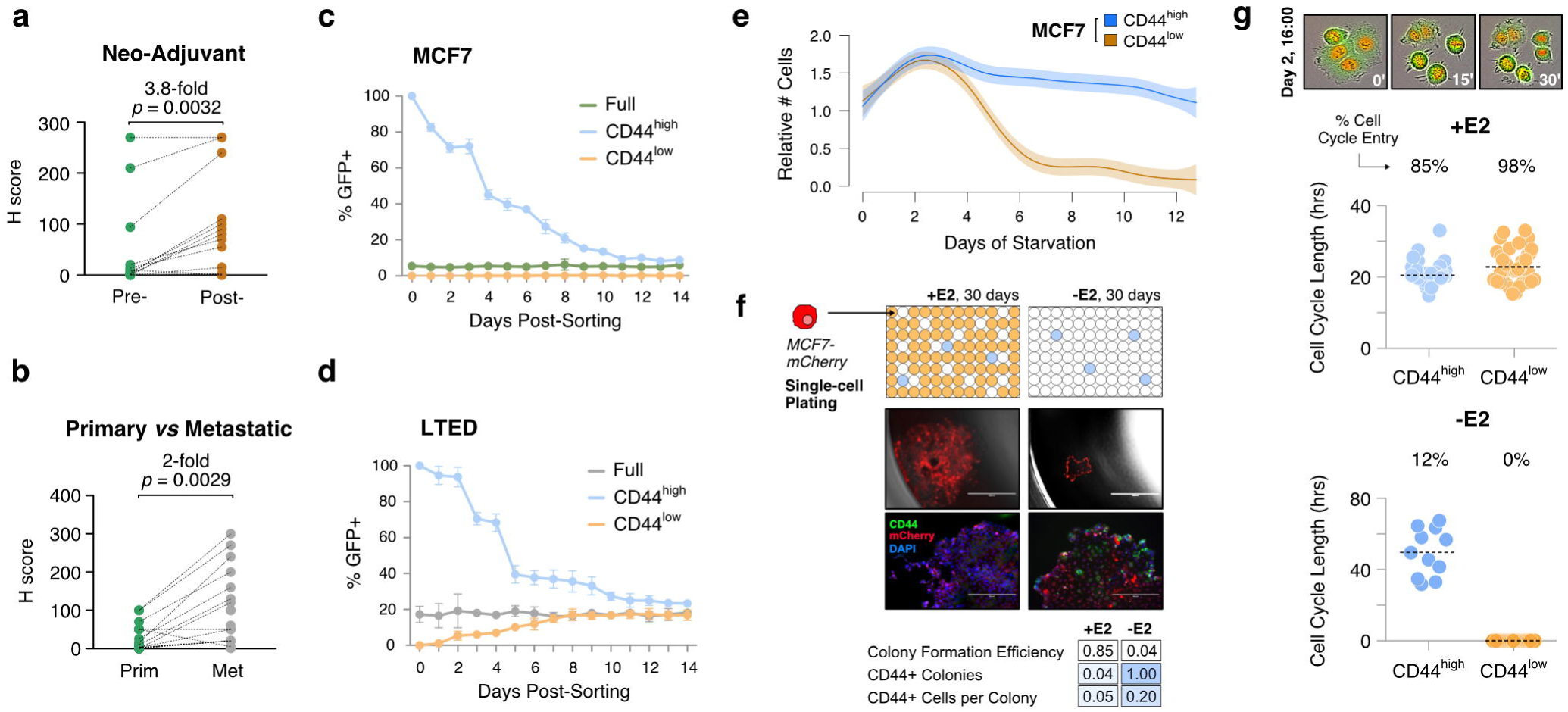
Characterisation of a phenotypically distinct population of plastic cells in luminal breast cancer. **(a)** CD44 expression in neo-adjuvant AI patients (pre- and post- treatment; *p*-value from two-tailed paired *t*-test). **(b)** Same as (a), but in matched AI-treated primary-metastatic (*p*-value from Wilcoxon signed-rank test). **(c)** Reconstitution experiments from FACS-sorted MCF7-CD44^GFP-high^ or MCF7- CD44^GFP-low^ cells. **(d)** Same as (c) but using FACS-sorted LTED cells. **(e)** Survival curves of MCF7-CD44^GFP-high^ and MCF7-CD44^GFP-low^ cells in oestrogen-deprived conditions. **(f)** Single-cell plating experiments in oestrogen-supplemented (+E2) or deprived conditions (-E2) for 30 days. From top to bottom: (i) schematic representation of the results; (ii) representative pictures of single wells after 30 days (scale bar = 400 μm); (iii) Immunofluorescence staining highlighting CD44 expression (scale bar = 200 μm); (iv) summary statistics. **(g)** Cell-cycle dynamics of MCF7-CD44^GFP-high^ and MCF7-CD44^GFP-low^ cells inferred from time-lapse imaging. The length of the cell cycle and percentage of cell entering the cell cycle are indicated for both oestrogen-supplemented (+E2; top) or deprived conditions (-E2; bottom).

To further investigate the plasticity of CD44^high^ cells *in vitro* at the single-cell level, we generated MCF7 and LTED cell lines with a GFP reporter expressed under the promoter of the CD44 gene (Supplementary Fig. 4). Reconstitution experiments from FACS-sorted cells showed that CD44^GFP-high^ cells could recapitulate all the functional aspects of endogenous CD44^high^ cells including cellular plasticity (Figure 2c). Interestingly, both CD44^GFP-high^ and CD44^GFP-low^ showed features of plasticity in fully resistant cells (Fig. 2d and Supplementary Fig. 3g). When MCF7 were challenged with short-term ET, only CD44^GFP-high^ cells appeared to adapt to it, while CD44^GFP-low^ cells were rapidly cleared out between days 4 and 7 (Fig. 2e). Single-cell plating experiments confirmed that only CD44^high^ cells could drive the formation of early colonies under E2-deprivation, but the colonies were significantly smaller compared to E2-supplemented conditions (Fig. 2f). These observations indicate combined cytostatic and cytotoxic effects of ET and that those cells that could adapt to the therapy originate within the CD44^high^ compartment. Extrapolation of cell-cycle dynamics of CD44^GFP-high^ and CD44^GFP-low^ cells from time-lapse imaging data revealed comparable cell-cycle length in E2-supplemented condition (Fig. 2g; +E2). Nevertheless, CD44^GFP-high^ cells had a significantly lower proportion of cells engaged in productive cell cycle entry, suggesting the existence of a low-proliferative subpopulation within the CD44^high^ compartment even under permissive environments. Under E2-deprivation, the CD44^GFP-low^ completely failed to undergo cell-cycle entry, while 12% of CD44^GFP-high^ managed to do one or more cell cycles, with a much longer latency (Fig. 2g; -E2).

Taken together, these results further support the idea that at least some of the cells in the CD44^GFP-high^ (but not CD44^GFP-low^) compartment have an increased ability to survive the acute phase of ET, and this correlates with their features of plasticity. This led us to hypothesise that non-genetic, transcriptional variability would reflect pre-existent, rare subpopulations in treatment-naïve cells with higher chances to survive and give rise to fully-resistant cells.

### Single-cell transcriptomic profiling unveils further heterogeneity of plastic cells

To investigate the transcriptional variability of CD44^high^ cells, we carried out sorting- driven, scRNA-seq of CD44-GFP luminal breast cancer cells. About 10,000 single cells in E2-supplemented condition were profiled (CD44^GFP-high^ and CD44^GFP-low^ in equal proportions; Fig 3a; in the remainder of the text, these two sorted subpopulations will be referred to as CD44^high^ and CD44^low^). Dimensionality reduction (Fig. 3a) highlighted a surprising similarity between the profiles of CD44^high^ and CD44^low^, except for a small percentage (∼4%) of CD44^high^ cells significantly departing from the main cluster. In line with this, differential expression analysis of the two subpopulations resulted in 10-fold less differentially regulated genes (DEGs) than those observed by comparing them to LTED (Fig. 3b; Supplementary Table 3). Nevertheless, CD44^high^ showed an overall, significantly higher transcriptomic variability (*p*-value < 2.2e-16; Wilcoxon rank-sum test) than CD44^low^ (Fig 3c).

**Figure 3.**
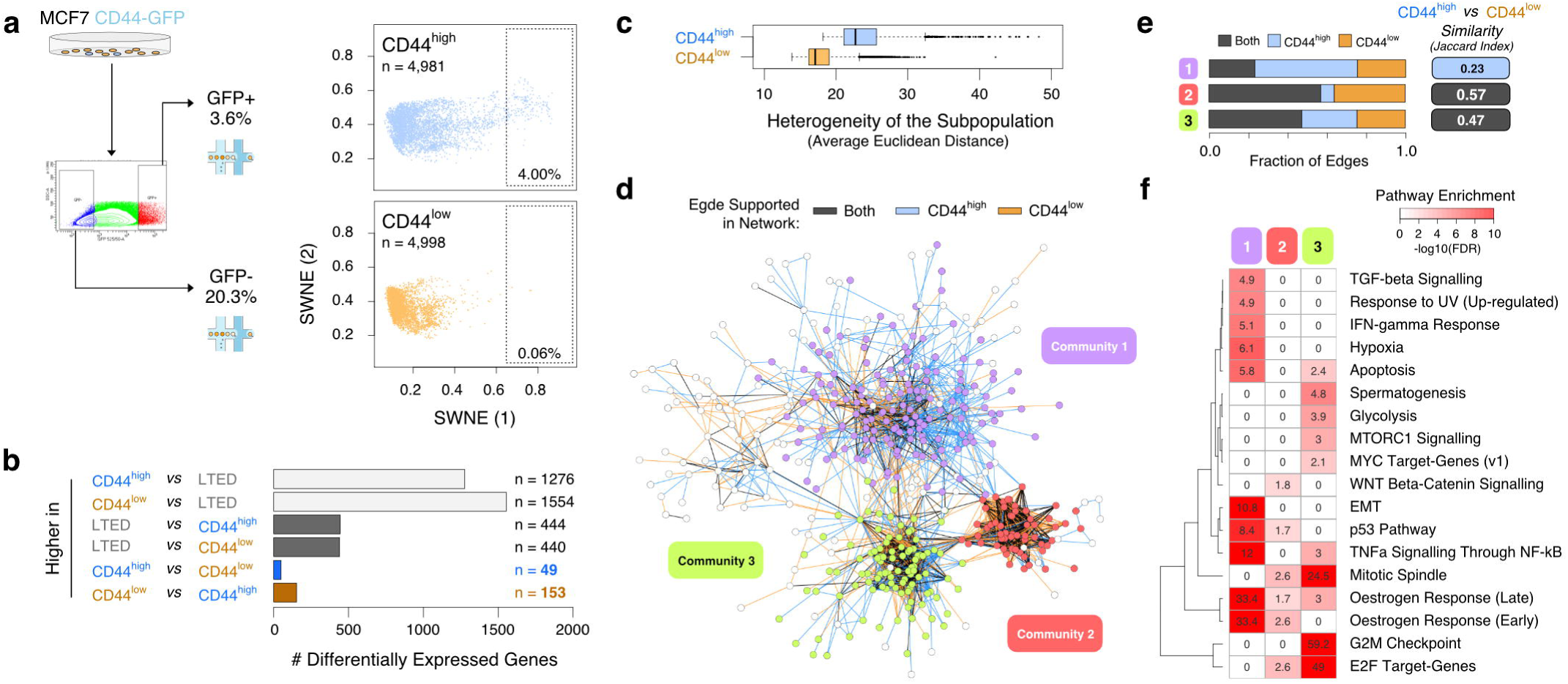
Single-cell transcriptomic profiling unveils further heterogeneity of plastic cells. **(a)** Schematic representation of the strategy to FACS-sorted MCF7- CD44^GFP-high^ (CD44^high^) and MCF7-CD44^GFP-low^ (CD44^low^) cells (left) along with results of dimensionality reduction for single-cell transcriptomes (right) (SWNE, *k* = 22); percentage of extreme outliers in the two subpopulations indicated in the bottom right corner. **(b)** Number of up-regulated genes in the indicated comparisons (FDR <= 0.05; AUC >= 0.6). **(c)** Cell-cell heterogeneity within CD44^high^ and CD44^low^ subpopulations. **(d)** Regulatory networks reconstructed using either CD44^high^ and CD44^low^ profiles were superimposed and the edges color-coded according to whether each edge was identified only in the CD44^high^ (blue), the CD44^low^ (orange) or both (dark grey) networks. Nodes in the three larger communities were color-coded accordingly. **(e)** Fraction of edges identified in the CD44^high^, the CD44^low^ or both networks, for each one of the three communities shown in (d). Similarity between CD44^high^ and CD44^low^ networks is also shown. **(f)** Enrichment analysis using the hallmark gene sets^38^ across the three communities shown in (d).

We next sought to systematically address whether the observed variability was the result of either an increased transcriptional noise specific to CD44^high^ cells (compatible with a bet-hedging mechanism) or instead the reflection of a regulated network (leading to coordinated expression of multiple genes in the same cell). We applied PIDC^37^, an algorithm using partial information decomposition (PID), to identify regulatory relationships between genes, and reconstructed the gene regulatory networks (GRNs) from the scRNA-seq profiles of CD44^high^ and CD44^low^ cells, separately (Supplementary Table 4). The two networks were merged and analysed to identify major communities (Fig. 3d). The larger of the three identified communities (#1 in Figs. 3d-f) showed low similarity between the CD44^high^ and CD44^low^ GRNs, with the majority of edges supported only by the CD44^high^ GRN (Fig. 3e). Pathway enrichment analyses^38^ for the genes in this community showed highly significant enrichments for oestrogen response, TNFα signalling, epithelial-mesenchymal transition and p53 pathway (Fig. 3f; *q*-value < 1e-8). These results strongly suggest that the variability specific to the CD44^high^ compartment is the result of the coordinated regulation of genes within well-defined subpopulations. With this in mind, we hypothesised a central role for these rare cells in the early phases of acute oestrogen deprivation (termed acute-ET).

### Single-cell profiling of acute response to endocrine therapy identifies a subpopulation of pre-adapted cells

To investigate the role of transcriptomic variability of plastic cells during acute-ET, we performed scRNA-seq experiments upon oestrogen deprivation (Supplementary Table 2). Continuous single-cell imaging suggested that cells within the CD44^low^ subpopulation started being differentially affected by acute-ET after 48 hours of treatment (Fig. 2e). We thus profiled gene expression data of about 10,000 single cells at 48 hours of starvation (Fig. 4a). Applying a stringent threshold on the first SWNE component, we could define a rare, pre-adapted (PA) subpopulation among plastic cells (CD44^high^) expressing a signature of acute-ET even in permissive E2- supplemented condition. The identification of PA cells was confirmed using an approach based on Random Forests classification (Fig. 4b, Supplementary Table 5 and Methods). Of note, PA cells are genetically indistinguishable from the other CD44^high^ cells and have not yet acquired any of the genetic re-arrangements of the fully resistant, LTED cells (Fig. 4c). Considering both approaches and both a lenient and a stringent threshold, PA cells are estimated to constitute 0.76 to 4% of the CD44^high^ cells, which correspond to 0.03 to 0.14% of the total MCF7 population. Overall, these data suggest that PA cells might represent the first step in the process of adaptation to acute-ET.

**Figure 4.**
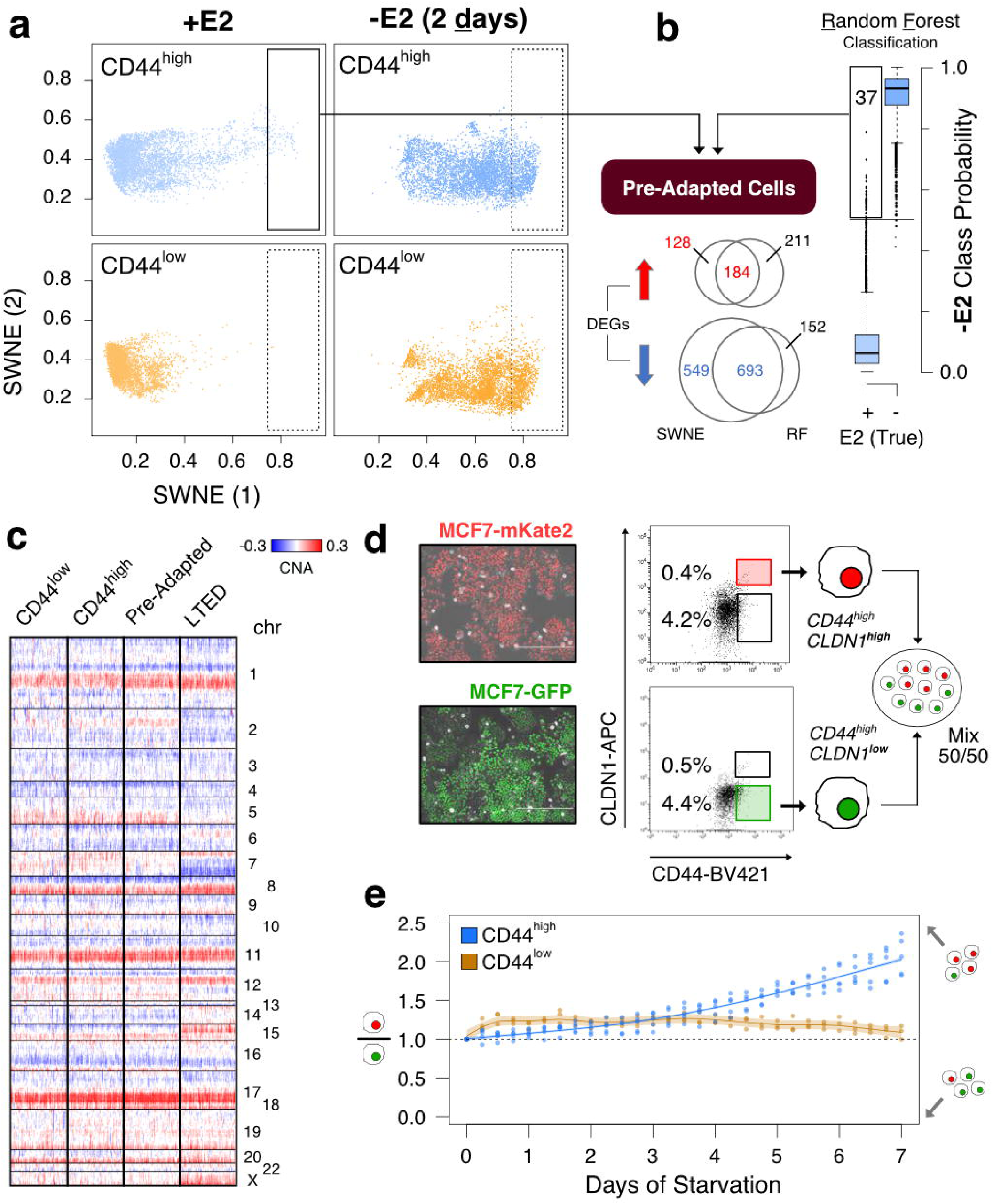
Single-cell profiling of acute response to endocrine therapy identifies a subpopulation of pre-adapted cells. **(a)** Dimensionality reduction of single-cell transcriptional profiles of oestrogen-supplemented (+E2; top) or deprived (-E2; 2 days) cells. Pre-adapted (PA) cells highlighted in boxes. **(b)** PA cells identification using two different strategies (SWNE and Random Forests). DEGs = Differentially Expressed Genes. **(c)** Copy number profiles of PA cells (n = 81), along with the same number of LTED, CD44^low^ and CD44^high^ (not PA) cells, as estimated from scRNA-seq profiles. **(d)** FACS-sorted PA cells (CD44^high^ and CLDN1^high^) stably labelled with mKate2 were mixed with other plastic cells (CD44^high^ and CLDN1^low^) stably labelled with GFP and followed up for 7 days upon oestrogen-deprivation **(e)**.

We then sought to validate if the PA transcriptional state would confer a survival advantage compared to other plastic cells exposed to acute-ET. First of all, we identified the *Claudin-1* gene (CLDN1) as a marker for PA cells (in combination with CD44). We then generated MCF7 cells stably labelled with either a nuclear GFP or mKate2 and leveraged this tool to follow two subpopulations over time after mixing them. The same amount of sorted PA cells (CD44^high^ and CLDN1^high^) was mixed with other plastic cells (CD44^high^ and CLDN1^low^; Fig. 4d). CD44^high^ CLDN1^high^ PA cells showed increased survival to acute-ET compared to CD44^high^ CLDN1^low^, with this effect increasing over time. As a control, no difference was observed between CLDN1^high^ and CLDN1^low^ from the CD44^low^ compartment. These data strongly support the hypothesis that PA cells have distinctive survival advantage under acute-ET (Fig. 4e).

We then further characterised these cells functionally. We focused on a stringent set of differentially expressed genes between the PA cells and the rest of the CD44^high^ cells in +E2 condition (312 up-regulated and 341 down-regulated; Figs. 4a,b and Supplementary Table 5). PA cells displayed features of mixed epithelial and mesenchymal traits, along with up-regulation of p53 pathway, cell polarity (apical junction components), and hypoxia (Fig. 5a, upper panel). PA cells also showed reduced ERα activity and down-regulation of the cell cycle machinery, while still expressing ESR1 (Fig. 5a, lower panel and Supplementary Fig. 5). Interestingly, both plastic and non-plastic cells lied on a continuum showing negative correlation between the expression of the genes of the cell cycle and of those in the signature of PA cells (Fig. 5b; Spearman’s Rank Correlation coefficient = −0.519; *p*-value < 2.2e- 16), with PA cells found at the edge of this spectrum. We finally sought to quantify the overlap between the PA cells signature (up-regulated genes) with the CD44^high^- enriched GRN we previously identified (Figs. 3d-f). Indeed, when we further dissected the GRN (community #1) into its two main components, we found extensive overlap between one of these components and the PA signature (Fig. 5c; *p*-value = 2.7e-21; hypergeometric test). This further supports the idea that the genes in this signature are part of a co-regulated network.

**Figure 5.**
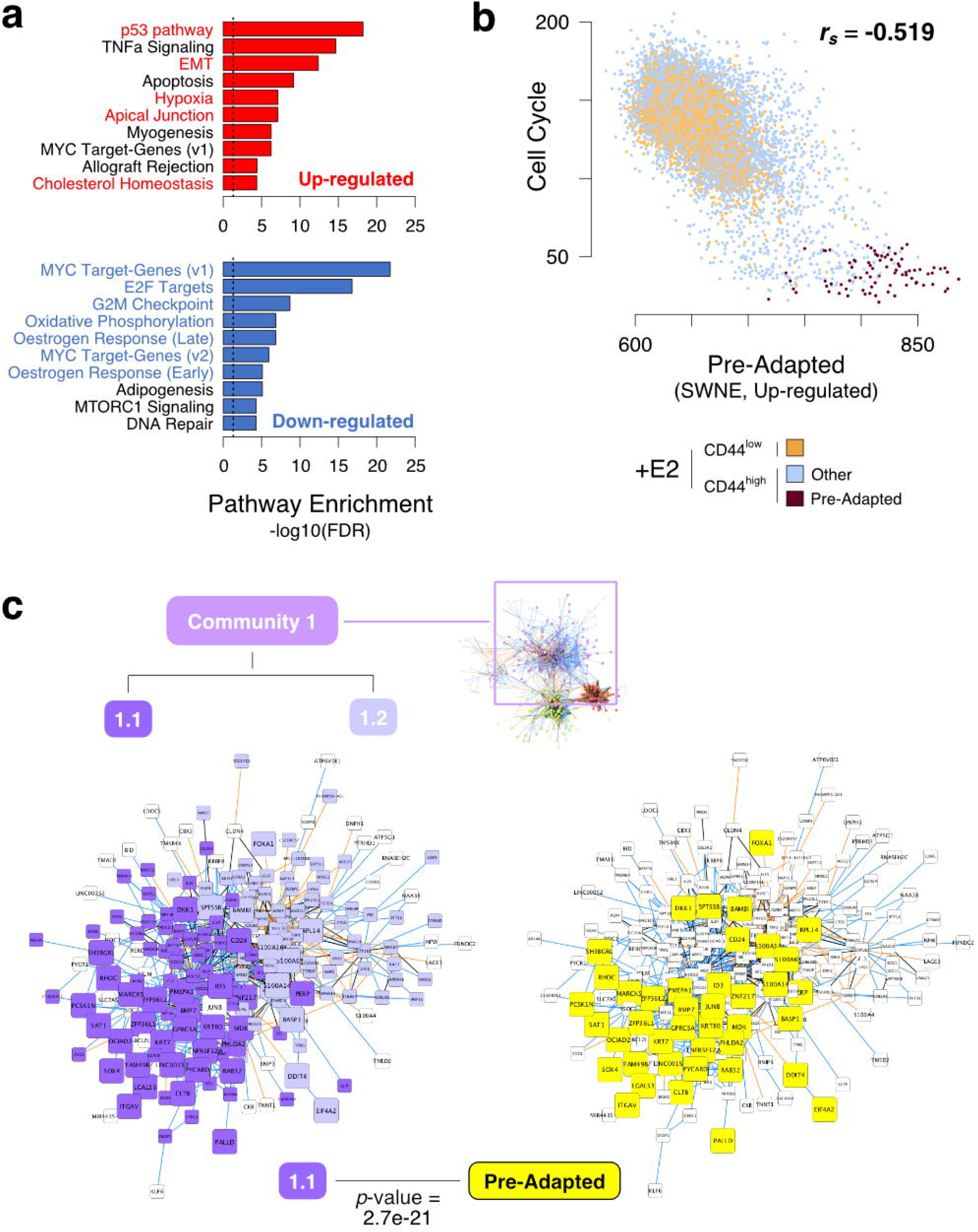
Functional characterisation of the signature of pre-adapted cells. **(a)** Hallmark gene sets^38^ enriched in genes either up- or down-regulated in PA cells. **(b)** Correlation analysis between expression of cell cycle marker genes and genes belonging to the PA-signature (up-regulation) at the single-cell level. rs = Spearman’s Rank Correlation Coefficient. **(c)** Same representation as in Fig. 3d but limited to community 1. Two sub-communities were identified (left) with community 1.1 being more strongly enriched for genes in the PA-signature (p-value from Hypergeometric Test).

Overall, these data support the hypothesis that plastic cells are non-genetically heterogeneous, and among them rare cells in the PA state have a survival advantage during acute-ET.

### Pre-adaptation is a persistent feature of acute-ET but differs from a signature of full resistance

While these analyses support a pivotal role for the PA phenotype in conferring a survival advantage during acute-ET, PA cells are still genetically indistinguishable from the rest of the cells. This suggests these cells do not represent the final step of drug resistance. Nevertheless, we aimed at determining whether longer exposure to acute-ET correlates with the persistence of the PA signature, and/or this also coincides with other reprogramming events. In order to capture the different dynamics of survival of CD44^high^ and CD44^low^ (Figs. 2e and 6a), we generated scRNA-seq profiles at 4 and 7 days of oestrogen deprivation (Supplementary Table 2), a period in which the relative number of CD44^high^ cells does not change while CD44^low^ undergo rapid extinction. Dimensionality reduction of >28k cells showed increased prevalence of the PA signature with time of starvation (Fig. 6a). Formal quantification using AUCell^39^ confirmed this trend (Fig. 6b, left panel and Fig. 6d). The same analysis using a LTED-specific signature (Methods and Fig. 1) failed to identify any cell expressing it during acute-ET (Fig. 6b, right panel). In line with this, the critical transcriptional pathways driving full resistance (i.e. cholesterol biosynthesis and re-activation of ERα signalling) were completely abrogated in PA and cells exposed to acute-ET (Figs. 6c,d). On the other hand, some of the pathways associated to PA phenotype (partial-EMT, cell polarity, hypoxia) were found to consistently increase during treatment.

**Figure 6.**
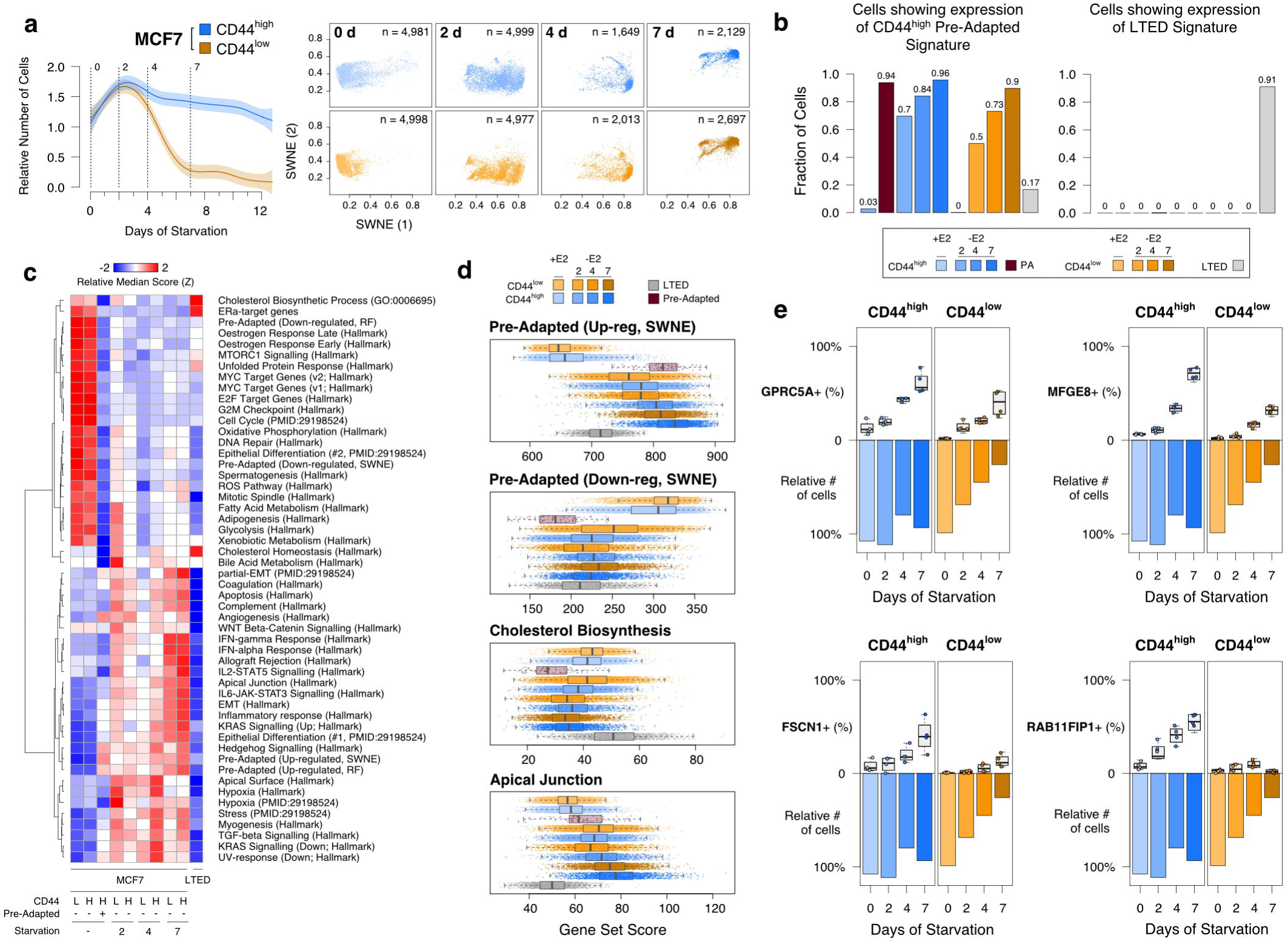
Pre-adaptation is a persistent feature of acute-ET but differs from a signature of full resistance. **(a)** Sampling design along with dimensionality reduction of single-cell transcriptional profiles of oestrogen-supplemented (day 0) or deprived (days 2, 4 and 7). **(b)** AUCell^39^ quantification of the fraction of single-cells showing transcriptome compatible with either the PA (left) or the LTED (right) signatures. **(c)** Selected gene set enrichment across all conditions profiled in this study. **(d)** Score distributions across cells. **(e)** Multi-marker tracing profiles for selected genes (box plots) in CD44^high^ and CD44^low^ cells upon oestrogen-deprivation. Survival (as relative number of residual cells) is also shown (bar plots).

Unexpectedly, while imaging showed that after 7 days >75% of the CD44^low^ died and were destined to extinction (Fig. 2f), the profiled CD44^high^ and CD44^low^ cells converged on the same transcriptional changes. We reasoned that since scRNA-seq experiments capture viable cells exclusively, we profiled only those cells that were still alive at day 7. Thus, we hypothesised that the PA-like transcriptional program is an intermediate bottleneck during acute-ET. In line with this, we discovered that CD44^low^ cells can occasionally up-regulate a signature overlapping that of PA cells, but this happens with lower efficiency (Fig. 6b) and it is not sufficient to give them the survival advantage shown by CD44^high^. To validate these observations at the protein level, we performed a multi-marker tracing profile, exploiting some marker genes (namely, GPRC5A, MFGE8, FSCN1 and RAB11FIP1) showing a trend of up-regulation with starvation time. This trend was confirmed at the protein level, with values consistently higher in CD44^high^ compared to CD44^low^ cells (Fig. 6e and Supplementary Fig. 6 and 7). Nevertheless, this did not prevent cells in the CD44^low^ compartment to die at an almost linear rate (Fig. 6e).

**Figure 7.**
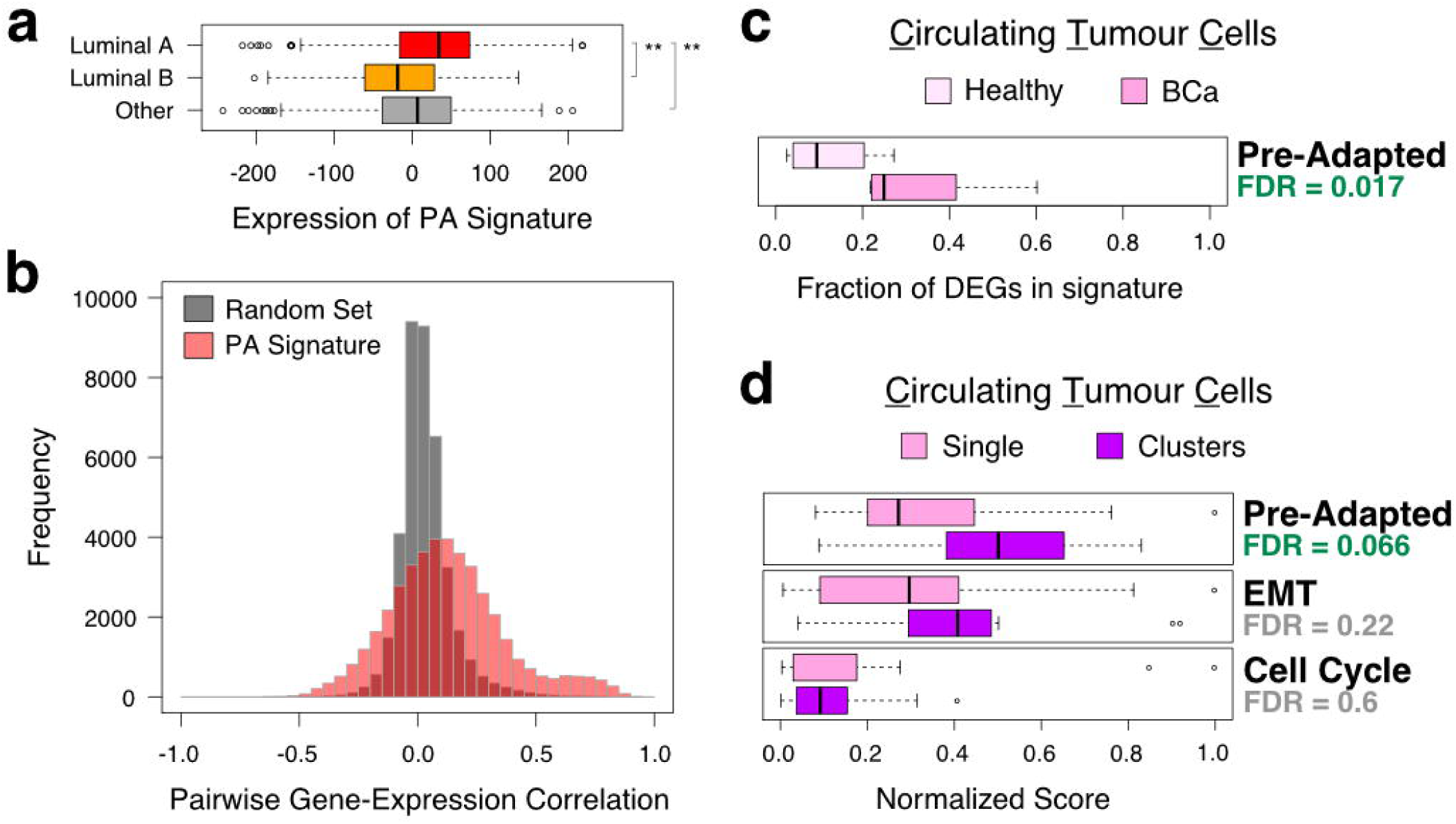
The pre-adapted signature is active in primary tumors and enriched in clusters of circulating tumor cells. **(a)** Expression of the PA signature in bulk RNA-seq samples from TCGA. **(b)** Distributions of Spearman’s rank correlation coefficients between expression profiles of genes in the PA signature (red) or a set of randomly picked genes of the same size (grey), across bulk RNA-seq samples from TCGA classified as luminal A. **(c)** Expression of the PA signature in circulating tumour cells (CTCs) compared to blood specimens from healthy donors. **(d)** Expression of the PA, the EMT and a cell cycle signatures in clusters of CTCs compared to single CTCs. False discovery rates estimated by permutations.

Taken together, these observations confirmed that the PA transcriptome is strongly selected by acute-ET. Nevertheless, the observed rapid expansion (Fig. 6b) seems incompatible with the strict selection of a pre-existing population^30^. The observation that also CD44^low^ cells can adopt a similar transcriptional profile in response to ET (despite being unable to survive) suggests the PA program is required but not sufficient to explain the survival of plastic cells to acute-ET (see Discussion).

### The pre-adapted signature is active in primary tumors and enriched in clusters of circulating tumor cells

We first looked for evidence of expression and co-regulation of genes up-regulated in PA cells, in 825 primary luminal breast tumours^40^. Tumours classified as luminal A showed significantly higher expression of the signature compared to luminal B (*p*- value < 2.2e-16; Wilcoxon rank-sum test) and TNBC/HER2+ lesions (*p*-value = 1.9e-8) (Fig. 7a). Of note, luminal A exhibit the longest latencies in relapse development amongst all BCa^41-43^. Considering >600 luminal A samples, we then checked the distribution of pairwise correlations between the expression pattern of the genes in the signature, as a proxy for co-regulation. Compared to a size-matched set of randomly picked genes, those in the PA signature showed significantly higher coefficients (Fig. 7b; *p*-value < 2.2e-16; Wilcoxon rank-sum test), with hundreds of pairs with values over 0.5 (Spearman’s Rank Correlation coefficient). These results further corroborate our previous observations that these genes tend to be controlled by the same GRNs and showed a trend of higher expression in luminal tumours with longer latency of recurrence (A vs B; Fig. 7a).

Given that some of the key pathways active in PA cells hinted to mixed epithelial and mesenchymal features, as well as cell polarity and migration, we asked if the PA phenotype could play a role in metastatic progression. Previous data strongly suggest that epithelial-like cluster of circulating tumour cells (CTCs) are responsible for 85-92% of metastatic dissemination^44^, with individual CTC showing more mesenchymal features playing a more limited role^45^. Interestingly, the PA signature was found significantly enriched in CTCs^45^ (Fig. 7c; *q*-value = 0.017, permutation test) and at even higher levels in clusters of CTCs^44^ (Fig. 7d; *q*-value = 0.066, permutation test). These results provide a further link between drug-induced adaptation and metastatic invasion^27,46^.

## Discussion

In this study, we leveraged an *in vitro* model to investigate the contribution of genetic and transcriptional heterogeneity to the development of resistance to ET in luminal breast cancer. As opposed to previous observations in melanoma, TNBC, lung and colorectal cancers, in which targeted therapy lead to the rapid emergence of fully-resistant cells^10–12,18,26^, we could not find any genetic or phenotypic clone showing features of resistance in treatment-naïve cells (Fig. 1). The same observation held true even after thoroughly dissecting the heterogeneity of the cells showing features of plasticity (Figs. 2-3). On the other hand, we could identify and characterise a small subpopulation (∼0.1% of the treatment-naïve cells) showing a pre-adapted (PA) phenotype (Fig. 4). These cells showed a 2-fold increased survival to acute-ET compared to other plastic cells, while non-plastic cells undergo complete extinction under selective pressure (Fig. 4e), along with mixed epithelial and mesenchymal features, and quiescence. Interestingly, while any cell (also those with no feature of plasticity) can adopt a transcriptional program overlapping that of the PA cells, only plastic cells can withstand acute-ET (Fig. 6a and 6e), with PA cells showing a more pronounced survival advantage (Fig. 4e). Finally, we found an enrichment of the PA signature in clusters of CTCs. To our knowledge, this is the first time a quiescent population from the primary tumour has been linked to both features of survival to therapy and of CTCs. Interestingly, it has been reported that early stage metastatic cells possess partial features of survival, dormancy and EMT, which all overlap with our PA signature^47^. A signature of partial-EMT has also been recently shown to be expressed in the cells at the leading edge of primary head and neck cancers^48^. It is tempting to speculate that PA cells might not only display a survival advantage during the early phases of the therapy but might also be the pioneers of micro-metastatic spread.

Surprisingly, we found that also cells with no features of plasticity were able to adopt the PA signature, even though with a much lower efficiency, which cannot prevent the extinction of the compartment after two weeks of oestrogen deprivation (Fig. 6). On top of this, 70% of the plastic cells adopted a PA signature within 48 hours of acute-ET (Fig. 6b). This fast transition to a diverse transcriptional state is hardly explained by straight Darwinian selection of a persister cell^30^. For reasons that remain to be investigated, plastic cells have a much higher probability than non-plastic ones to transition into a PA state, and this probability is dramatically increased by oestrogen deprivation. We reason that upon stress, plastic PA cells are better positioned than cells requiring transcriptional reprogramming, hence the observed difference in survival within the plastic compartment (Fig. 6e).

PA cells represent an obligated step toward the acquisition of resistance while still requiring substantial reprogramming to recapitulate features of full resistance cells. We propose that the delayed relapse common to ET treated patients might be mediated by similar processes, in which PA-like cells are selected for and stalled by ET for up to >10 years. This model would reconcile why ET are sometime effective for down staging neo-adjuvant patients but fail to clear micro-metastatic disease. Nevertheless, how this bottleneck affects the progression of the tumour requires further investigation. Future studies on the necessary steps and their timing of occurrence during treatment must be carried out in order to expose potential vulnerabilities of these quiescent cells.

## Methods

### Cell lines

Cells were originated as previously described^27^. MCF7 cells were maintained in Dulbecco’s modified Eagle’s medium (DMEM) containing 10% fetal calf serum (FCS). Long-term oestrogen-deprived cells (LTED) were derived from MCF7 after one year oestrogen deprivation as previously described and were maintained in phenol-red free DMEM containing 10% charcoal stripped fetal calf serum (SFCS)^27^. Both media were supplemented with 2 mM L-glutamine, 100 units/mL penicillin and streptomycin. 10^−8^ M estradiol (E2758 Sigma) was added routinely to MCF7. Primary metastatic breast cancer cells were derived from pleural effusions of patients with metastatic breast cancers. The pleural effusion (PE) cells were maintained in DMEM containing 10% fetal calf serum (FCS) and 2 mM L-glutamine, 100 units/mL penicillin and streptomycin.

### Plasmids

pLVX-IRES-mCherry-puro lentiviral vector (Cambridge Bioscience, Cambridge, UK) was used to infect MCF7 and LTED cells. CD44 reporter GFP cells were established with CD44CR1-IRES-GFP-puro lentiviral vector (Tebu-Bioscience). Stable and polyclonal cell populations were established after puromycin selection (0.5 μg/ml). NucLight Green lentivirus (IncuCyte, 4626) and NucLight Red Lentivirus (IncuCyte, 4627) were used to infect MCF7. Stable and polyclonal cell populations were established after Zeocin selection (300 μg/ml). H2B-mCherry-puro lentiviral vector was used to infect stable CD44 reporter GFP cells. Stable and polyclonal cell populations were established after sorting.

### Antibodies

Anti-ERα antibody (Vector Laboratories, VP-E613) 1:100 for immunofluorescence (IF) and anti-ERα (Santa Cruz, HC-20) 1:1000 for western blot (WB), anti-CD44 antibody (Santa Cruz, sc-7297,) 1:200 for IF and 1:100 for immunohistochemistry (IHC), anti-pan Cytochemistry antibody (Abcam, ab17154) 1:200 for IF, anti-FGFR4 antibody (Abcam, ab44971) 1:100 for IF, anti-FSCN1(Sigma, HPA005723) 1:100 for IF, anti-MFGE8 (Sigma, HPA002807) 1:100 for IF, anti-RAB11FIP1 (Sigma, HPA023904) 1:100 for IF, anti-GPRC5A (Sigma, HPA007928) 1:100 for IF, anti-caspase3 (Merk, MAB10753) 1:100 for IF.

### FACS analysis

Cells were cultured to 70 to 80% confluence and detached from the cell culture flasks using EDTA. Cell pellets were obtained and washed with cold phosphate-buffered saline (PBS) containing 1% FCS and 5mM EDTA. All further steps were performed on ice and all centrifugation steps at 4°C. Fluorochrome-conjugated monoclonal antibodies against human CD44 (FITC, BD Pharmingen; BV421, BD Pharmingen), Claudin1 (APC, R&D systems), and their isotype controls were added to the cell suspension at concentrations recommended by the manufacturer (BD Biosciences) and incubated at 4 °C in the dark for 30 min. The labelled cells and CD44 reporter GFP cells were washed in PBS and then were analyzed on a FACSAria (BD Biosciences). Gating was set to relevant isotype control (IgG-FITC)-labeled cells or unstained cells for each cell line. Propidium iodide (Bio-Rad, 1351101) and DRAQ7 (BioLegend, 424001) were used for the dead cell removal.

### Soft Agar Colony-Forming Assay

Anchorage independent cell growth was carried out in six-well tissue culture plates. A 1-mL layer of 0.6% agar (DIFCO Laboratories) in appropriate cell culture medium was solidified at the bottom of each well. Cells to be assayed were suspended in 1 mL of 0.3% agar in medium. 1 × 10^4^ cells were seeded in each dish. After 4 weeks of incubation at 37 °C in 5% CO2, colonies were visualized by staining with 0.02% Crystal violet.

### Mammosphere Culture

Cells were plated as single cells at a density of 5×10^2^ viable cells/well in ultralow attachment 6 well plates (Corning, CLS3814). Cells were grown in a serum-free DMEM or phenol-red free DMEM, supplemented with B27 (Invitrogen, 17504-044), 20 ng/mL EGF (Sigma, E9644) and 20 ng/mL bFGF (R&D systems, 233-FB-025). Mammospheres were grown for 10–14 d and phase contrast images were obtained using the ImageXpress Micro microscope (Molecular Devices). For the second-generation experiment, first-generation mammospheres were collected from multiple wells and spun at 500 × g per 5 minutes. The pellet was resuspended in 50μl Trypsin and the sample was passed 25 times through a sterile needle to get single cell suspension. The same density of cells as in first generation culture was seeded, and cells were allowed to grow for 14

### Immunofluorescence

Briefly, 10^4^ cells were seeded on chamber slides (Lab-Tek). On the final day, cells were washed twice with PBS at room temperature and 4% PFA/PBS was added for 15 minutes. Cells were washed twice with PBS and NH4Cl was added as quencher. 0.2% Triton/PBS was added for 5 minutes. 10% BSA/PBS was used as blocking reagent. 5% BSA/PBS was used to dilute primary antibodies and Alexa-fluor 488, 568, 594, 647 labelled anti-rabbit or anti-mouse secondary antibodies (ThermoFisher). Nuclei were counterstained with DAPI and were mounted in ProLong Antifade Mountant (ThermoFisher, P36941). Pictures were acquired using EVOS microscope system (Advanced Microscopy Group, Bothell, WA, USA) or Zeiss Axiovert 200M inverted microscope.

### Tissue microarray (TMA) of paired primary and secondary breast cancers

Twenty primary breast carcinomas with a paired metastasis were acquired from the pathology archives of Charing Cross Hospital, London, UK. A tissue microarray was constructed using a manual microarrayer and 0.6 mm punches. The tissue microarray was immunohistochemically profiled for CD44 (Santa Cruz sc-7297). Antigen retrieval was performed using 0.01M citrate buffer, pH 6.0 followed by blocking in 0.3% hydrogen peroxide in PBS, then in normal goat serum (20 μl per ml) for 30 min. The primary antibody was incubated overnight at 4°C at 1:100, and then detected using anti-mouse secondary antibody (Vector Laboratories), Vectastain Elite peroxidase ABC kit, and ImmPACT DAB kit (Vector Laboratories). Subsequently, 4μm TMA sections were immuno-stained using the optimized staining protocol, including negative controls (omission of the primary antibody). Staining was scored based on the H-score and Allred Quick score by three independent investigators (including one consultant pathologist) blinded to the clinicopathological characteristics of patients. H = (3 × % of strongly stained cells) + (2 × % of moderately stained cells) + (1 × % of weakly stained cells) + (0 × % of cells without staining). Negative controls were performed by omission of the primary antibody.

### Neo-adjuvant patient selection

All clinical data from patients operated at the European Institute of Oncology (IEO) were prospectively entered in an Institutional data base. For the present study we retrieved data from patients with a neoadjuvant treatment from 1999 to 2014. We selected patients having had pre-surgical biopsy and surgery in our Institute, in order to have their sample analyzed in the same laboratory. We randomly selected 20 patients with luminal tumors, treated by neoadjuvant hormone only (aromatase inhibitors), 10 responders and 10 non-responders.

### Reconstitution assay

After sorting with CD44 using FACS Aria III (BD Biosciences), 10^5^ cells of CD44^high^ and CD44^low^ were seeded on 6 well plates and incubated for 7 days. After 7 days, cells were trypsinized and stained with anti-CD44 antibody (FITC, BD) for FACS. After sorting with GFP using FACS Aria III (BD Biosciences), 10^5^ cells of CD44^GFP-high^ and CD44^GFP-low^ were seeded on 6 well plates with oestradiol supplement or deprivation. Five pictures per condition were taken using an EVOS microscope system (Advanced Microscopy Group, Bothell, WA, USA) during 14 days. Fifty different fields were counted. The % of GFP positive cells was calculated by number of GFP positive cells/ number of total cells x100.

### Monoclonal Assay

Using FACS Aria III (BD Biosciences), single cell of mCherry-MCF7 was seeded on 96 well plates with oestradiol supplement or deprivation. Single cell was confirmed using an EVOS microscope system (Advanced Microscopy Group, Bothell, WA, USA). The cells were incubated for 30 days, and colonies were counted on EVOS microscope system.

### Live cell imaging and Data Analysis

After sorting with GFP by FACS Aria III (BD Biosciences), 10^5^ cells of H2B mCherry MCF7 CD44^rep^ GFP^pos^ and GFP^neg^ were seeded on 6 well plates with estradiol supplement or deprivation. Time-lapse live cell imaging was performed on IncuCyte ZOOM (Essen BioScience) equipped with temperature, humidity and CO2 control. Images were acquired every 6 hours with 10X plan fluorescence objectives for a proliferation assay and every 15 min with 20x up to 10 days for a cell cycle analysis. Excitation (Ex) and emission (Em) filters sets (Chroma Technology Corporation) were as follows: CFP, 427-10nm (Ex), 483-32 nm (Em); YFP, 504-12 nm (Ex), 542-27 nm (Em); mCherry, 589-15 nm (Ex), 632-22 nm (Em). Micromanager 1.3 was used for acquisition of time-lapse images. All data analysis was done with scripts written in Matlab (Mathworks) or using Cell Profiler (Broad Institute) and ImageJ (National Institutes of Health). Symmetric/Asymmetric/conversion analyses were performed on a total of 200 cells. Each cell was monitored for the first 3 cell divisions (1 cell to 2 cells, 2 cells to 4 cells, 4 cells to eight cells). Symmetric division was scored if the daughter cell matched the mother. Asymmetric was scored if the daughter cell did not match the mother. Conversion was scored if cell changed CD44 status without cell division (at least 4 hours pre-or post- division). Cell cycle speed was established by calculating the time intervening between two consecutive metaphase plates.

### Statistical analyses

Unless specified otherwise, all the analyses and plots were performed in the statistical computing environment R v3 (www.r-project.org).

### Single cells preparation

Single cells were prepared from a full population of MCF7 and LTED, or from sorting MCF7 CD44-GFP reporter cells by the level of GFP expression at different time points of E2 deprivation. After centrifugation, single cells were washed with PBS and were re-suspended with a buffer (Ca^++^/Mg^++^ free PBS + 0.04% BSA) at 1,000 cells/µl.

### Single-cell RNA sequencing

Viability was confirmed to be > 90% in all samples using acridine orange/propidium iodide dye with LUNA-FL Dual Fluorescence Cell Counter (Logos Biosystems, L20001). Single cell suspensions were loaded on a Chromium Single Cell 3’ Chip (10X Genomics) and were run in the Chromium Controller to generate single-cell gel bead-in-emulsions using the 10x genomics 3’ Chromium v2.0 platform as per manufacturer’s instructions. Single-cell RNA-seq libraries were prepared according to the manufacturer’s protocol and the library quality was confirmed with a Bioanalyzer High-Sensitivity DNA Kit (Agilent, 5067- 4627) and a Qubit dsDNA HS Assay Kit (ThermoFisher, Q32851). Samples were pooled up to four and sequenced on an Illumina HiSeq 4000 according to standard 10X Genomics protocol.

### Single-cell RNA-seq raw data analysis

cellRanger (v2.1.1) was run on the raw data using GRCh38 annotation (v1.2.0). Output from cellRanger was loaded into R using the function *load_cellranger_matrix_h5* from package *cellranger* (v1.1.0; genome = “GRCh38”). Datasets were merged according to gene names. All cells sampled were retained except for flow-sorted CD44^high^ and CD44^low^ either in +E2 media or starved for two days, for which the top 5,000 cells in terms of UMIs per cell were considered. In order to robustly detect transcriptional states, a recent paper suggested to consider a coverage of at least 1,500 detected genes per cell^49^. A filter on cells showing at least 1,500 detected genes per cell and at least 5,000 UMIs per cell was then applied. After that, reads mapping on mitochondrial genes were excluded. Before normalization, a series of filtering steps were performed. To do that, data were imported in *Seurat* (v2.3.4)^50^ and scaled (*NormalizeData* function using normalization.method = “LogNormalize”, scale.factor = 10000, followed by the *ScaleData* function). A filtering step was then performed based on the cumulative level of expression (the sum of the Seurat-scaled values) of three housekeeping genes (GAPDH, RPL26 and RPL36)^51^. Manual inspection of these values versus the number of UMIs per cell (or the number of genes with non-zero expression per cell) revealed no correlation between the two. Nevertheless, a number of cells showed very low expression for these genes. Cells showing housekeeping gene expression in the bottom 1% were then excluded from further analyses. At last, genes expressed in less than 20 cells were excluded. Across-cells normalization was performed using the R package *Scran* (v1.6.9)^52^. Raw counts were imported into a SCE object using the *newSCESet* function; size factors were calculated using *computeSumFactors* (sizes = seq(20, 250, 10)), on data pre-clustered through *quickCluster*.

### Estimation of Copy number alterations (CNAs) from scRNA-seq data

CNAs were estimated directly from the scRNA-seq data, using an approach similar to the one used by Patel at al.^53^. Only genes expressed in >=25 cells were considered.

A reference gene expression profile was generated based on published scRNA-seq profiles of hormone-responsive luminal cells (termed L2)^54^, using only the datasets obtained using a droplet-based approach. After normalizing each single-cell profile based on the total number of detected transcripts to a fixed constant (10,000), a pseudo-bulk profile for the L2-cells was derived using the mean expression value of each gene across all cells.

Before running the actual CNAs quantification, all the raw scRNA-seq datasets generated in this study (after filtering, pre-normalization) and the pseudo-bulk profile generated as described above were linearly normalized to a constant (10,000) and log-scaled (pseudo-count set to 1).

First of all, chromosomal coordinates of all genes were retrieved using the *biomaRt* R package (v2.34.2; host set to “jul2015.archive.ensembl.org”)^55^. This way, genes were sorted by chromosomal coordinates. A genome-wide scan was then conducted using a sliding window of 100 genes, with a step of 10. Using the *rollapply* function from the zoo package in R (v1.8-3), mean value of expression in each bin was calculated for each single cell, as well as for the reference profile. The resulting genome-wide profile from each single cell was then linearly regressed against the reference estimate (using the function *lm*). The residuals were then considered as a proxy for CNAs and plotted in the form of heat maps. Single-cell, CNAs profiles were hierarchically clustered (hclust, method="ward.D2”) and shown as a circular dendrogram using *circlize_dendrogram* from R package dendextend (v1.8.0). In case of MCF7 and LTED (full populations), CNAs were estimated on all the cells. In case of the identified pre-adapted cells, the same number of cells (n = 81) was randomly sampled from the other groups of cells.

### Estimation of CNAs from ChIP-input-DNA

Reads were aligned to the hg19 human reference genome using bowtie2 (v2.3.4.3)^56^. Aligned reads were converted to BAM files, sorted and indexed using Samtools (v1.9)^57^. Duplicated reads were marked and removed using Picard *MarkDuplicates* (v2.1.1; REMOVE_DUPLICATES=true). Only uniquely mapped reads were retained for further analyses. Copy numbers were inferred using CNVkit tools (v.0.9.4.dev0)^58^, as described here: https://cnvkit.readthedocs.io/en/stable/pipeline.html. CNVkit was run with the default parameters of the *batch* command after creating a flat reference genome as suggested in the manual using the command *reference*.

### Dimensionality reduction and clustering

Normalized data was then imported in *Seurat* and scaled. Variable genes were identified using the *FindVariableGenes* function (mean.function = ExpMean, dispersion.function = LogVMR, x.low.cutoff = 0.01, x.high.cutoff = 6, y.cutoff = 0.01, num.bin=100). Principal component analysis (PCA) was run using variable genes as input, and the top 50 components were kept. Clusters were then identified using *FindClusters* (resolution = 0.6). Considering only those variable genes identified as described above (Similarity Weighted Nonnegative Embedding (SWNE)^32^ was applied to further reduce the dimensionality of the data. The *k* parameter was estimated using *FindNumFactors* on a subsample of 1,000 cells (loss = “mse”, 2 to 50 as range of values, with a step of 2). The choice of *k* is determined by randomly set 20% of the gene expression matrix as missing, followed by finding the factorization that best imputes the missing values, minimizing the mean squared error. Using this parameter, non-negative matrix factorization was then run through *RunNMF* (alpha = 0, init = “ica”, loss = “mse”), followed by *EmbedSWNE* (alpha.exp = 1.25, snn.exp = 1.0, n_pull = n_4, dist.use = “IC”). For this step, the shared nearest neighbour (SNN) matrix calculated by the *FindClusters* function of *Seurat* was used.

### Differential expression analysis

The Two-sample Likelihood Ratio Test implemented in the *LRT* function of the *MAST* R package (v1.4.1)^59^ was used to identify marker genes for a given sample or cluster. Briefly, each cell was either flagged as either belonging to the sample (or the cluster) or not. Those genes identified as up-regulated in the cluster at FDR <= 0.05 (Benjamini-Hochberg correction)^60^ and showing an Area Under the Curve (AUC) >= 0.6 were classified as markers for the sample or the cluster. The AUC is an estimate on how accurately a certain gene predicts a cell as part of a certain sample or cluster. AUCs were calculated using the *ROCR* R package^61^.

### Functional enrichment analyses

For functional enrichment analyses a selected number of gene sets was employed. The fifty Hallmark gene sets from the Molecular Signature Database (MSigDB)^38^ were downloaded from the MSigDB website October 19, 2017. Gene sets from Puram et al.^48^ (Table S7), along with a manually curated list of ERα-core target genes (BYSL, GREB1, HEY2, MPHOSPH10, MYB, NIP7, RARA, SLC9A3R1, TFF1, XBP1) were also considered. For a given subset of cells, each gene set was scored separately as the sum of the normalized expression values of all the genes in the set. The resulting distributions were then used for statistical testing and visualization.

### Single cell gene regulatory network inference

Networks were inferred separately for CD44^high^ and CD44^low^ cells, with nodes representing genes and edges representing statistical dependencies between gene pairs. For each dataset, genes expressed in fewer than 20% of cells were excluded; then all possible network edges were ranked using the PIDC network inference algorithm^37^ implemented in NetworkInference.jl (http://github.com/Tchanders/NetworkInference.jl), with expression data for each gene discretized independently into 6 bins of equal width; finally, a network was defined keeping the 2,000 highest ranking edges. The two networks were then superimposed to form an overlapping network with edges belonging (i) only to the CD44^high^ network, (ii) only to the CD44^low^ network, or (iii) to both networks. Communities were detected in the overlapping network (and recursively in each community) using the label propagation method implemented in LightGraphs.jl (http://github.com/JuliaGraphs/LightGraphs.jl). Communities were required to include at least 10 nodes. Similarity of the CD44^high^ and CD44^low^ networks within each community was calculated using the Jaccard index: the number of edges in the community that belong to both the CD44^high^ and CD44^low^ networks divided by the total number of edges in the community; an edge was deemed to belong to a community if it connected two nodes in the community.

### Identification of the pre-adapted cells

Two different strategies were employed to identify the pre-adapted cells. The first one takes advantage of SWNE; a threshold was applied on the first component and the cells showing extreme values (>= 0.75) were labeled as pre-adapted. The second strategy leverages random forests classifiers^62^. First of all, the datasets of CD44^high^ cells in +E2 media and starved conditions (2 days) were split into training and testing sets, using 10% and 90% of the cells, respectively. The training set was then used to call the DEGs between the two conditions (+E2 *vs* starved), using the procedure described in the *Differential expression analysis* paragraph above. These DEGs were used as input features to train a random forest classifier, using the *randomForest* R package (v4.6-14; default parameters). This model was then used to test the remaining data. Those cells in the testing set labelled as +E2 that were showing a probability >50% of being classified as starved were considered pre-adapted.

AUCell^39^ (R package v1.0.0) was the used to quantify the activity of the pre-adapted signatures (and of other signatures, whenever indicated in the text) in single cells. First of all, normalized data was processed using the *AUCell_buildRankings* function. The resulting rankings, along with the signatures of interest, were then subject to function *AUCell_calcAUC* (aucMaxRank set to 5% of the number of input genes). Following inspection of the resulting distributions, thresholds were then manually set to 0.37, 0.18 and 0.32 for the signatures of pre-adapted cells either based on SWNE or random forests, or for the LTED signature (defined as those genes up-regulated in LTED *vs* MCF7, as described in the *Differential expression analysis* paragraph above).

### Re-analysis of published primary samples

Bulk RNA-seq datasets for 1,222 breast cancer samples were downloaded from the GDC (Genomic Data Commons)^40^ data portal (http://portal.gdc.cancer.gov/) using *gdc-client*, according to metadata obtained on July 25, 2018. Gene features were normalized to sequencing depth. Given that only a fraction of the samples was pre-classified using PAM50^63^, *k*- nearest neighbors (*k*-NN) classification was employed to impute the rest of the samples. This was performed via the *knn* function in the R package class (v7.3-14), using the pre-classified samples as training data. Unclassified samples were ascribed to a particular subtype only when showing >60% probability of being assigned to that class. Spearman’s correlations between expression profiles of pairs of genes were calculated on the depth-normalized values. Prior to calculating signature scores, these numbers were further log2-transformed (pseudo-count set to 1) and scaled to *z*-score gene-wise.

### Re-analysis of published datasets for circulating tumour cells (CTCs) and clusters of CTCs

Normalized data for CTCs collected at five time points from a single patient along with identically processed blood specimens from 10 healthy donors^45^ were downloaded from GEO (GSE41245). For each capture, the log2-fold-change between EPCAM+ cells and the matched IgG+ cells (control) was calculated. DEGs were defined as those genes showing a linear fold-change between EPCAM+ cells and control >= 1.5. The fraction of DEGs overlapping the genes in the pre-adapted signature was then calculated for each pair. To test if the observed difference between the fraction of DEGs in CTCs and in healthy specimens was random, a P-value was calculated using the Wilcoxon rank-sum test. The corresponding false discovery rate (FDR) was estimated by 1,000 permutations.

Raw data for individual CTC-clusters (median of 3 cells per cluster) and numerically matched pools of single CTCs from the same specimen^44^ were downloaded from GEO (GSE51827). Each profile was normalized by depth, then a profile-specific score was derived for the signature of the pre-adapted cells by summing the normalized expression values of all genes in the signature. These numbers were then divided by the maximum across all profiles. To test if the observed difference between the values obtained for the clusters against the matched pools of CTCs, a P-value was calculated using the Wilcoxon rank-sum test. The corresponding false discovery rate (FDR) was estimated by 1,000 permutations.

### GFP+ cells quantification

A custom Python script (available on request) was employed to segment images based on DAPI (to count the total number of cells) and GFP signal (to quantify the fraction of GFP+ cells).

## Supporting information

## Data availability

Raw sequencing data was deposited at the Gene Expression Omnibus (GEO) under accession number GSE119693. Reviewers can access the data using token mzgzmeowzfctneb and following the instructions provided at this link: http://www.ncbi.nlm.nih.gov/geo/query/acc.cgi?acc=GSE122743

## Author contribution

SP.H., I.B., and L.M. planned the research; SP.H. and Y.L. performed the experiments; I.B., SP.H., T.C., K.M., and L.M. analysed the data; G.C., N.R., G.P., S.A. and R.C. contributed technical support; SP.H., I.B., and L.M. wrote the manuscript.

## Acknowledgments

We want to acknowledge and thanks all patients and their families for the support and for donating the research samples. The authors gratefully acknowledge infrastructure support from the Cancer Research UK Imperial Centre, the Imperial Experimental Cancer Medicine Centre and the National Institute for Health Research Imperial Biomedical Research Centre. L.M. was supported by a CRUK fellowship (P64250) and Imperial Junior Fellowship (G53019). SP.H. was supported by a Basic Science Research Program through the National Research Foundation of Korea (NRF-2013R1A1A1011832) and by a CRUK programme award (CRUK C37/A18784). I.B. was supported by CRUK funding (P64250) and by an Imperial College Research Fellowship. The Imperial College Healthcare NHS Trust Tissue Bank provided tissue samples. Consent was collected at IEO and Imperial College (Project R15036) by the respective Tissue Banks. Other investigators may have received samples from these same tissues. The views expressed are those of the author(s) and not necessarily those of the NHS, the NIHR or the Department of Health.

---

**Figure S1.**
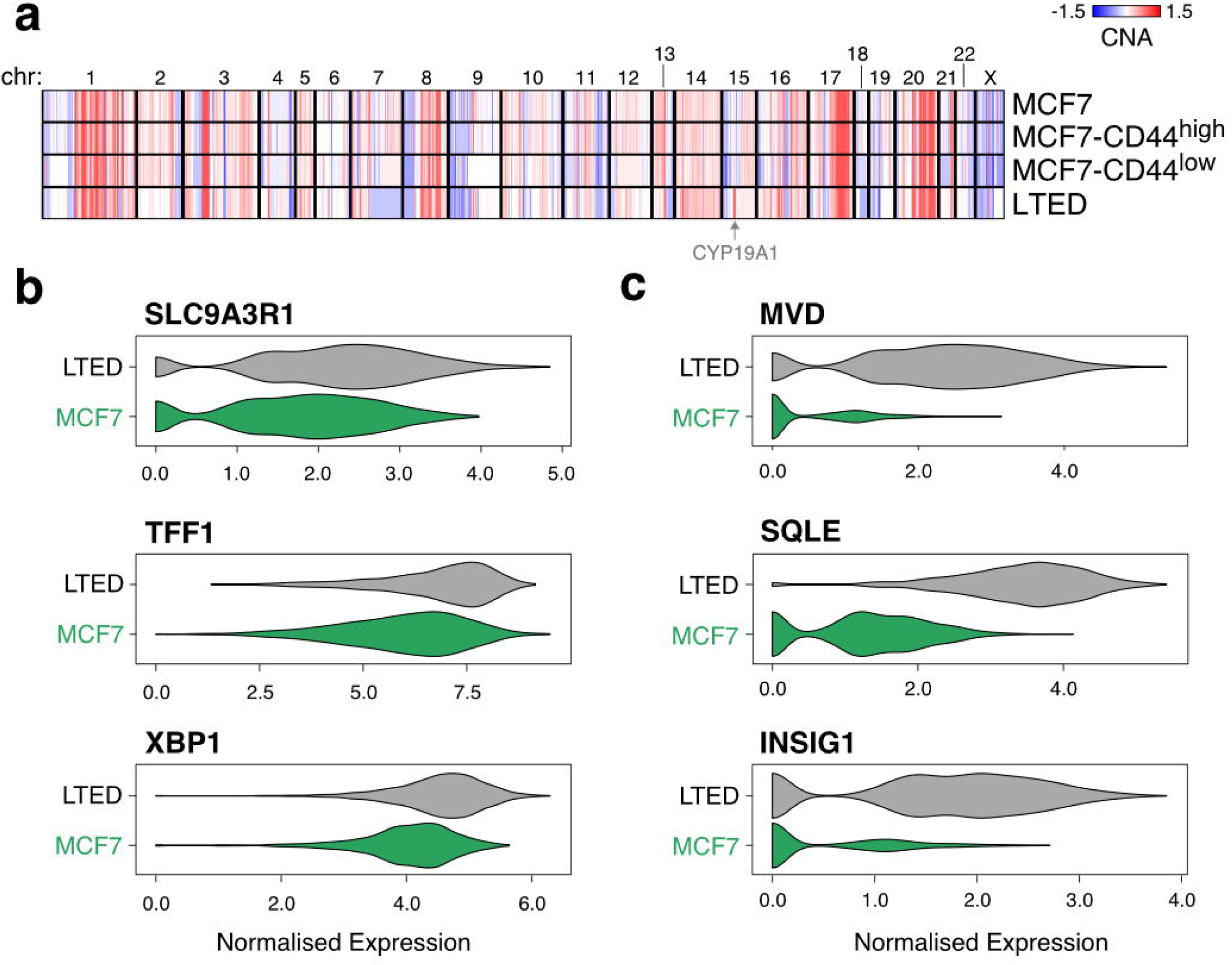
**(a)** Copy number profiles of the MCF7 (full population), MCF7-CD44^high^, MCF7-CD44^low^ and LTED, as estimated from ChIP-input DNAs. **(b-c)** Normalized expression of a selection of ER-target genes (b) and genes involved in cholesterol biosynthesis and homeostasis (c) in single MCF7 and LTED cells.

**Figure S2.**
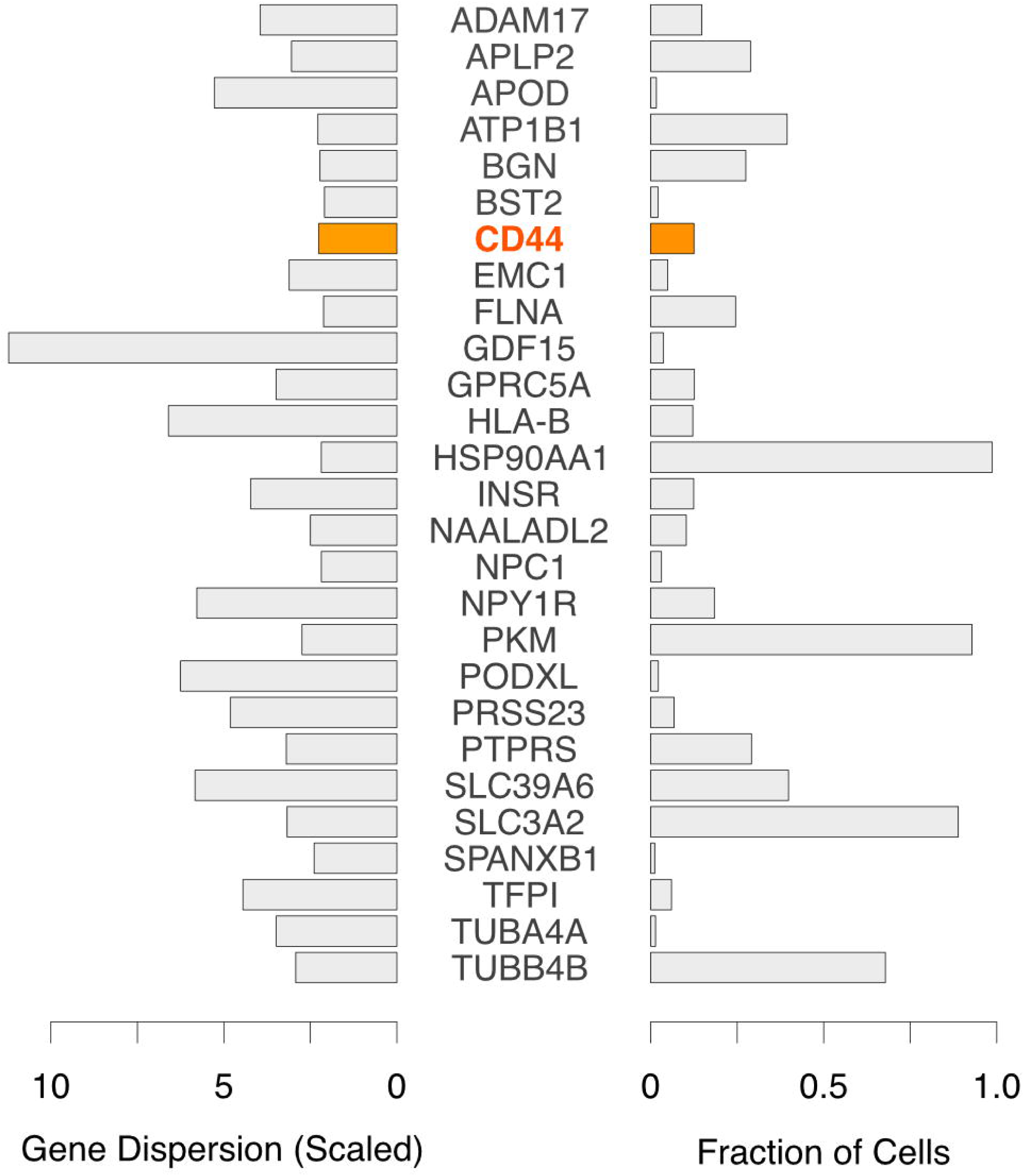
**(a)** Heterogeneously Expressed Surface Markers as estimated from scRNA-seq profiling. Genes were selected as showing dispersion >= 2 (n = 27) out of Fraction of Cells >= 1% (N = 778).

**Figure S3.**
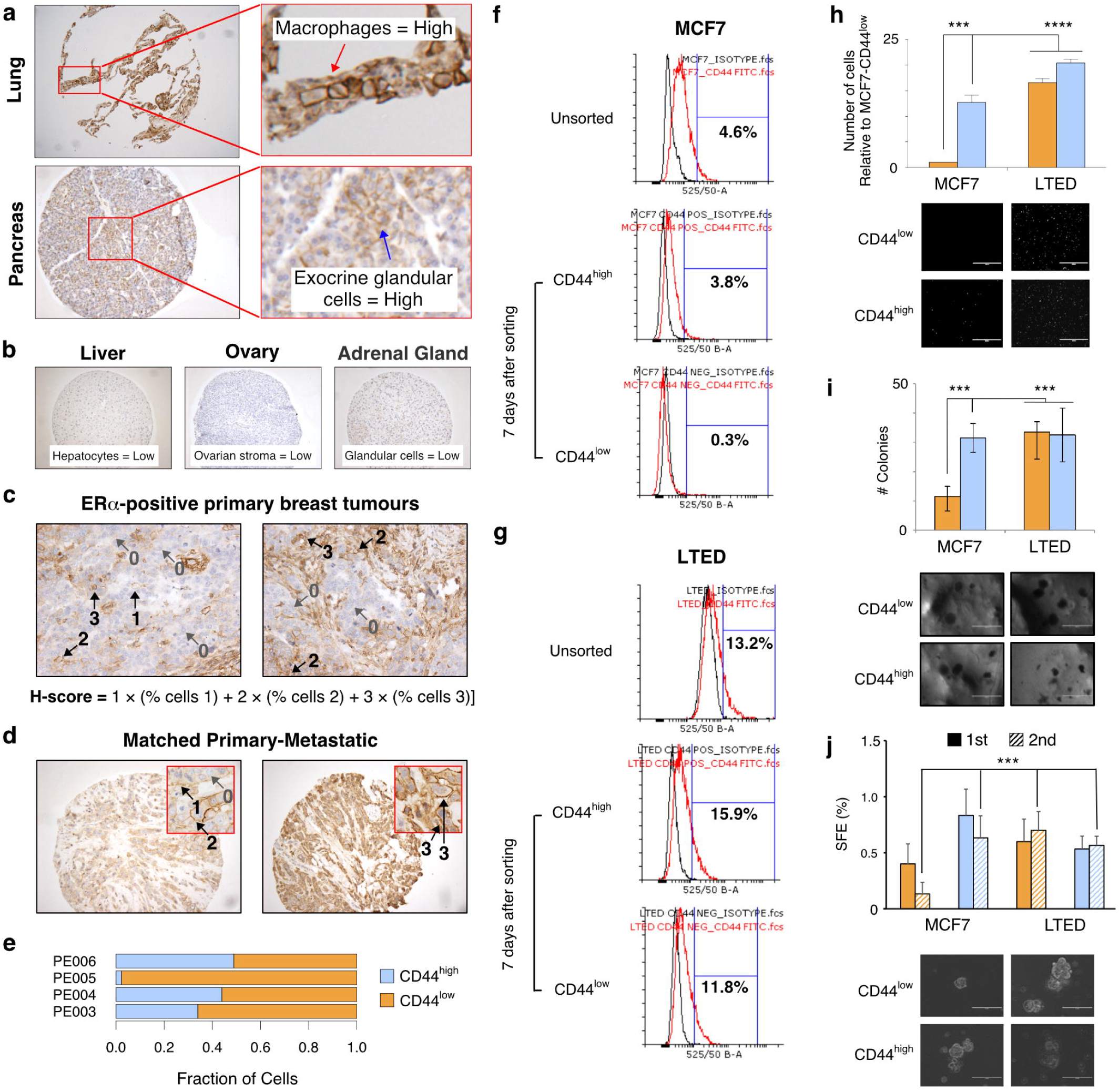
**(a-d)** Immunohistochemical staining of CD44 in two primary tumours (c) and in one matched primary-metastatic pair. Staining from lung and pancreas (a) and from liver, ovary and adrenal gland (b) are shown as positive and negative controls, respectively. **(e)** FACS quantification of CD44^high^ cells in pleural effusion cells from four patients. **(f-g)** FACS quantification of CD44^high^ cells in MCF7 (f) or LTED (g) cells. Unsorted populations were compared to purified CD44^high^ or CD44^low^ cells cultured for 7 days in oestrogen-supplemented medium. **(h-i)** Invasion (h) and colony formation (i) assays for CD44^high^ and CD44^low^ cells sorted from either MCF7 or LTED cells. Representative images shown below the plots (*p*-values estimated using two-tailed paired *t*-tests). **(j)** Mammosphere-forming efficiency (SFE) of for CD44^high^ and CD44^low^ cells sorted from either MCF7 or LTED cells. Representative images shown below the plots (*p*-values through ANOVA) Scale bars = 1,000 μm (i), 100 μm (j). Standard deviation of the mean estimated from 3 replicates is shown. * *p* <= 0.05, ** *p* <= 0.01, *** *p* <= 0.001, **** *p* <= 0.0001).

**Figure S4.**
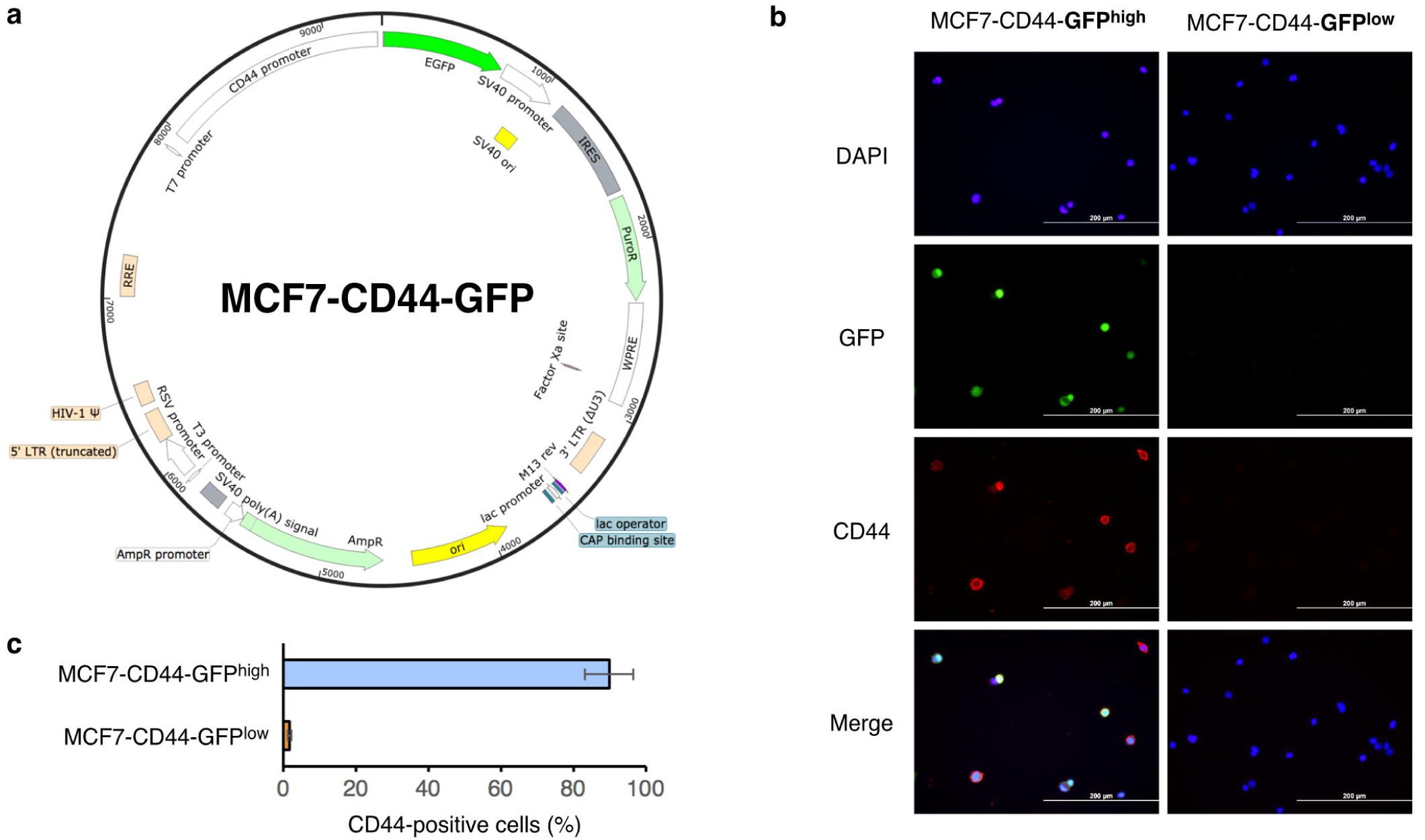
**(a)** Overview of the construct used to derive MCF7 stably expressing GFP under the control of the promoter of the CD44 gene. **(b)** Representative immunofluorescent images of FACS-sorted GFP^high^ and GFP^low^ cells, stained for DAPI and CD44. Scale bar = 200 μm. **(c)** Quantification of the fraction of FACS- sorted GFP^high^ and GFP^low^ cells from (b). Standard deviation of the mean estimated from 3 replicates is shown.

**Figure S5.**
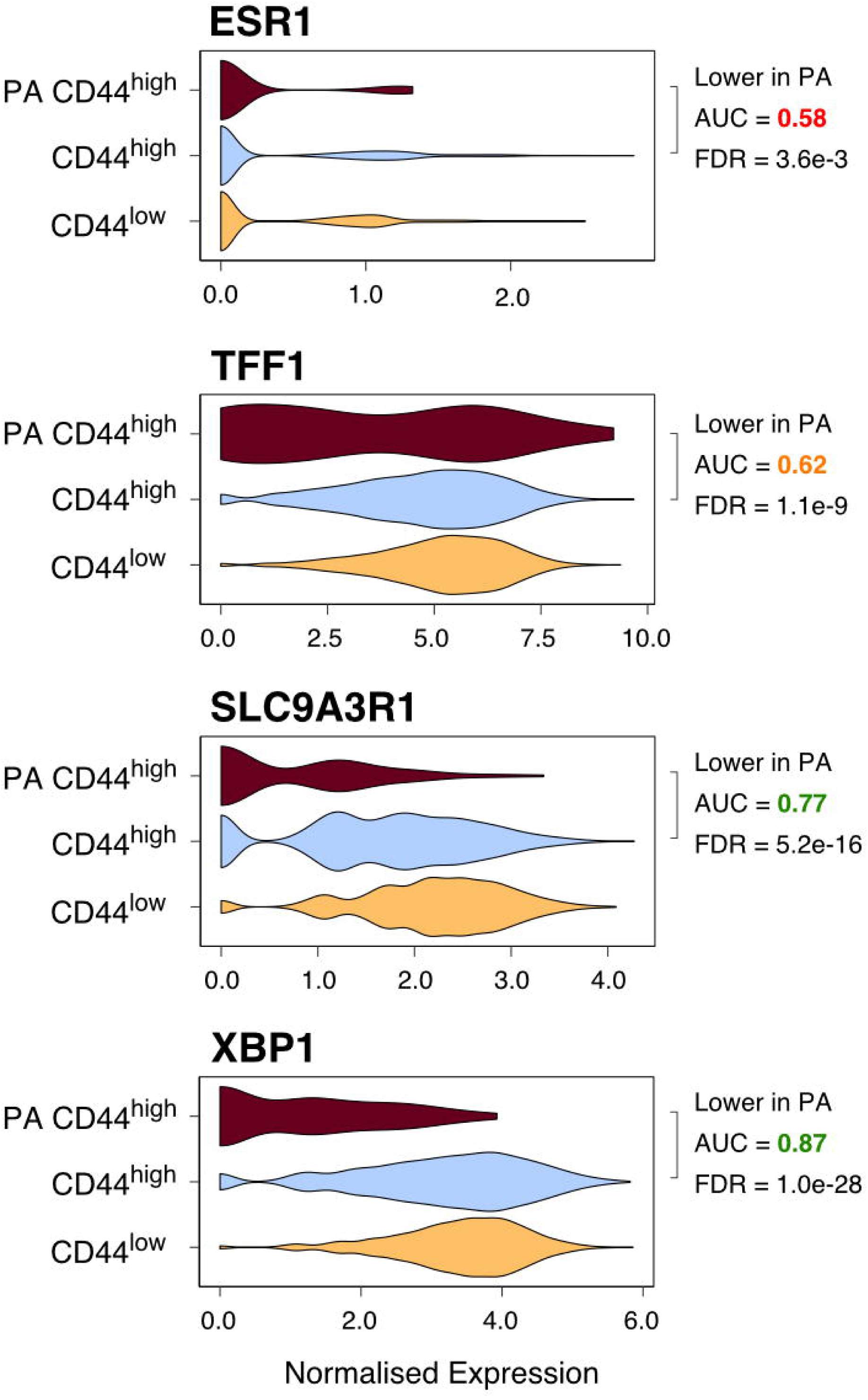
Normalized expression of ESR1 and of selected ER-target genes in the indicated subpopulations of MCF7 cells.

**Figure S6.**
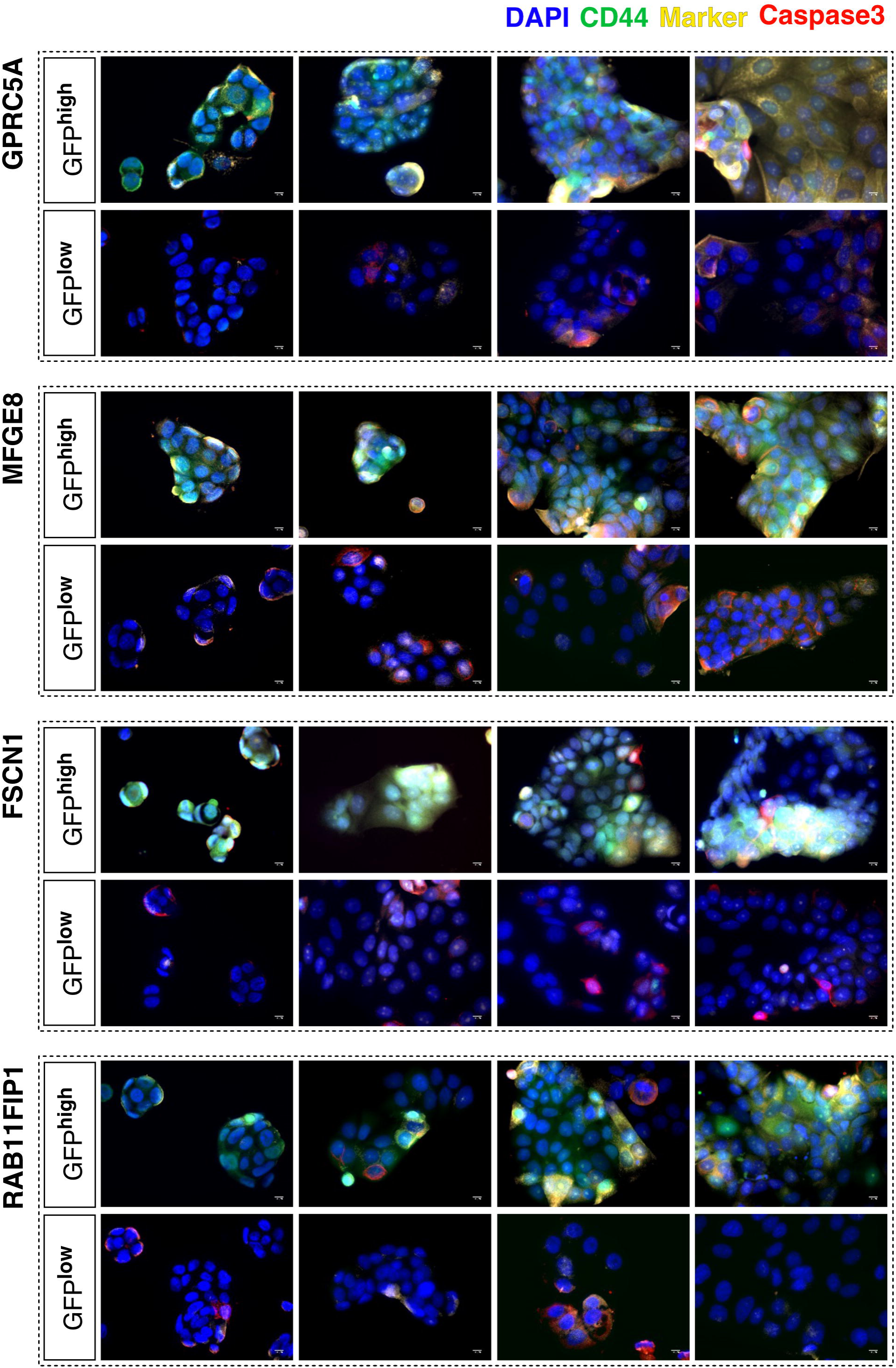
Representative images of purified CD44^high^ or CD44^low^ cells, stained for DAPI, CD44, Caspase3 and the indicated marker gene. Scale bar = 10 μm.

**Figure S7.**
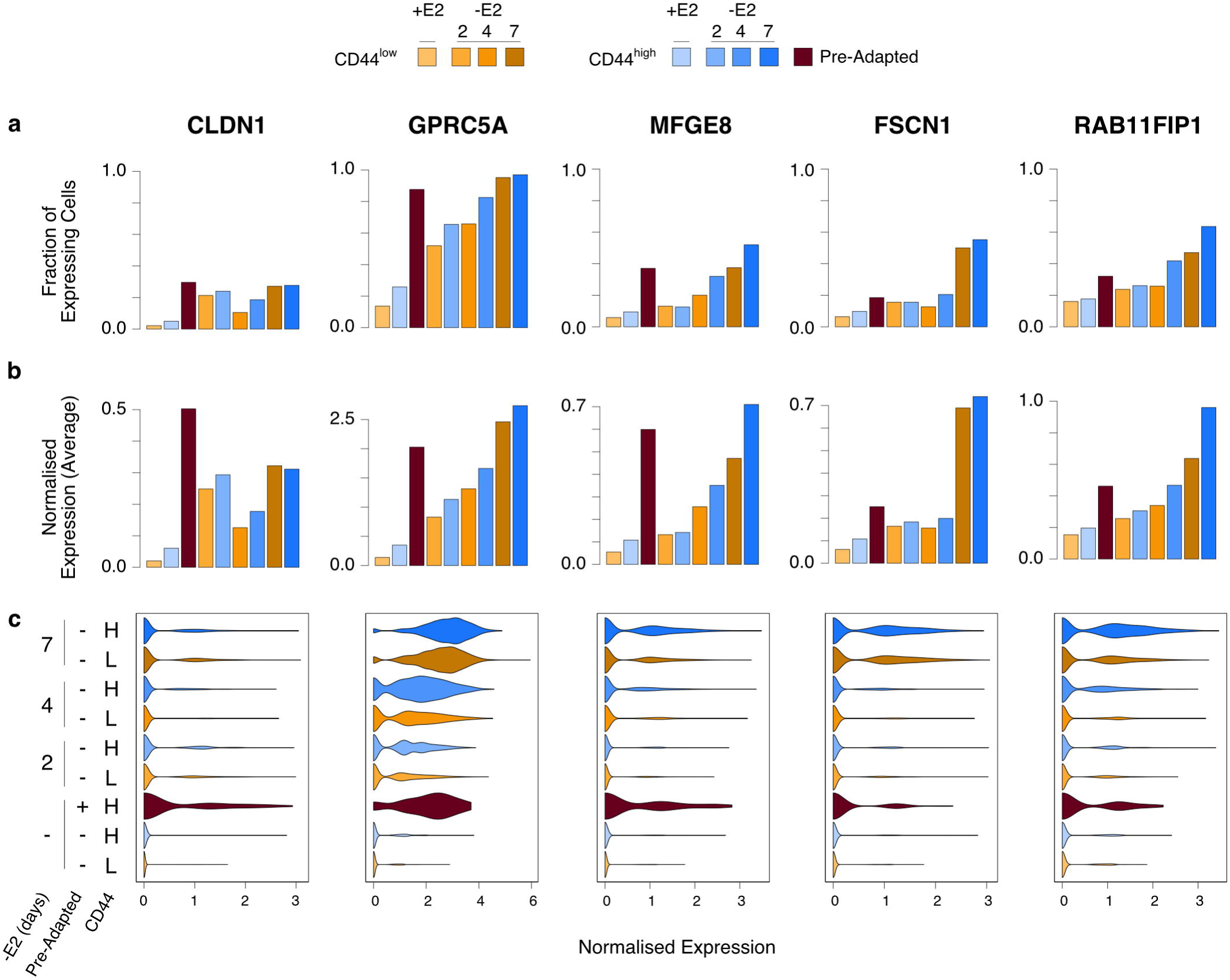
**(a-c)** For the indicated genes, the fraction of cells expressing them (a) along with the average (b) and the distribution (c) of normalized expression values are shown for the indicated subpopulations (top panel).

**Table S1.** Results of the re-analysis of treatment-naïve HR+/HER2- primary tumours from Razavi et al. 2018^31^.

**Table S2.** Summary statistics for all scRNA-seq samples reported in this study.

**Table S3.** Differentially expressed genes (DEGs) between MCF7-CD44^high^ (CD44H), MCF7-CD44^low^ (CD44L) and LTED. For each comparison, the false discovery rate (FDR) estimated using MAST^59^, along with the area under the curve (AUC) and the fraction of cells expressing each DEG in the compared subpopulations are indicated.

**Table S4.** Network reconstructed using PIDC^37^. For each inferred regulatory interaction, the two nodes are shown (genes 1 and 2), along with information indicating whether the edge is supported by either the network inferred from CD44^high^ and CD44^low^ data only, or both. For both genes, the table indicates the network component, whether the gene falls into one of the three main network communities (i.e. the three larger components) and into any of the two sub-communities of community 1, and whether the gene is part of the PA signature.

**Table S5.** Genes either up- or down-regulated in PA cells, using either the SWNE- based or the Random-Forest-based (RF) approach. For each list, the false discovery rate (FDR) estimated using MAST^59^, along with the area under the curve (AUC) and the fraction of cells expressing each DEG in the compared subpopulations are indicated.

## References

1. Early Breast Cancer Trialists’ Collaborative Group (EBCTCG), E. B. C. T. C. G. et al. Relevance of breast cancer hormone receptors and other factors to the efficacy of adjuvant tamoxifen: patient-level meta-analysis of randomised trials. Lancet (London, England) 378, 771–84 (2011).

2. Early Breast Cancer Trialists’ Collaborative Group (EBCTCG). Aromatase inhibitors versus tamoxifen in early breast cancer: patient-level meta-analysis of the randomised trials. Lancet (London, England) 386, 1341–1352 (2015).

3. Asselain, B. et al. Long-term outcomes for neoadjuvant versus adjuvant chemotherapy in early breast cancer: meta-analysis of individual patient data from ten randomised trials. Lancet Oncol. 19, 27–39 (2018).

4. Pan, H. et al. 20-Year Risks of Breast-Cancer Recurrence after Stopping Endocrine Therapy at 5 Years. N. Engl. J. Med. 377, 1836–1846 (2017).

5. Casasent, A. K. et al. Multiclonal Invasion in Breast Tumors Identified by Topographic Single Cell Sequencing. Cell 172, 205–217.e12 (2018).

6. Yap, T. A., Gerlinger, M., Futreal, P. A., Pusztai, L. & Swanton, C. Intratumor Heterogeneity: Seeing the Wood for the Trees. Sci. Transl. Med. 4, (2012).

7. Polyak, K. Heterogeneity in breast cancer. J. Clin. Invest. 121, 3786–8 (2011).

8. Turajlic, S. et al. Tracking Cancer Evolution Reveals Constrained Routes to Metastases: TRACERx Renal. Cell 173, 581–594.e12 (2018).

9. Jamal-Hanjani, M. et al. Tracking the Evolution of Non–Small-Cell Lung Cancer. N. Engl. J. Med. 376, 2109–2121 (2017).

10. Xue, Y. et al. An approach to suppress the evolution of resistance in BRAFV600E-mutant cancer. Nat. Med. 23, 929 (2017).

11. Nazarian, R. et al. Melanomas acquire resistance to B-RAF(V600E) inhibition by RTK or N-RAS upregulation. Nature 468, 973–977 (2010).

12. Misale, S. et al. Emergence of KRAS mutations and acquired resistance to anti-EGFR therapy in colorectal cancer. Nature 486, 532–536 (2012).

13. Johnson, B. E. et al. Mutational Analysis Reveals the Origin and Therapy-Driven Evolution of Recurrent Glioma. Science (80-.). 343, 189–193 (2014).

14. Robinson, D. R. et al. Activating ESR1 mutations in hormone-resistant metastatic breast cancer. Nat. Genet. 45, 1446–51 (2013).

15. Magnani, L. et al. Acquired CYP19A1 amplification is an early specific mechanism of aromatase inhibitor resistance in ERα metastatic breast cancer. Nat. Genet. 49, 444–450 (2017).

16. Siegel, M. B. et al. Integrated RNA and DNA sequencing reveals early drivers of metastatic breast cancer. J. Clin. Invest. 128, 1371–1383 (2018).

17. Clatot, F. et al. Kinetics, prognostic and predictive values of ESR1 circulating mutations in metastatic breast cancer patients progressing on aromatase inhibitor. Oncotarget 7, 74448–74459 (2016).

18. Kim, C. et al. Chemoresistance Evolution in Triple-Negative Breast Cancer Delineated by Single-Cell Sequencing. Cell 0, (2018).

19. Tirosh, I. et al. Dissecting the multicellular ecosystem of metastatic melanoma by single-cell RNA-seq. Science 352, 189–96 (2016).

20. Hinohara, K. et al. KDM5 Histone Demethylase Activity Links Cellular Transcriptomic Heterogeneity to Therapeutic Resistance. Cancer Cell (2018). doi:10.1016/J.CCELL.2018.10.014

21. Patten, D. K. et al. Enhancer mapping uncovers phenotypic heterogeneity and evolution in patients with luminal breast cancer. Nat. Med. 24, 1469–1480 (2018).

22. Rambow, F. et al. Toward Minimal Residual Disease-Directed Therapy in Melanoma. Cell 0, (2018).

23. Ebinger, S. et al. Characterization of Rare, Dormant, and Therapy-Resistant Cells in Acute Lymphoblastic Leukemia. Cancer Cell 30, 849–862 (2016).

24. Sharma, S. V. et al. A Chromatin-Mediated Reversible Drug-Tolerant State in Cancer Cell Subpopulations. Cell 141, 69–80 (2010).

25. Karaayvaz, M. et al. Unravelling subclonal heterogeneity and aggressive disease states in TNBC through single-cell RNA-seq. Nat. Commun. 9, 3588 (2018).

26. Shaffer, S. M. et al. Rare cell variability and drug-induced reprogramming as a mode of cancer drug resistance. Nature 546, 431–435 (2017).

27. Nguyen, V. T. M. et al. Differential epigenetic reprogramming in response to specific endocrine therapies promotes cholesterol biosynthesis and cellular invasion. Nat. Commun. 6, 10044 (2015).

28. Martin, L.-A. et al. Discovery of naturally occurring ESR1 mutations in breast cancer cell lines modelling endocrine resistance. Nat. Commun. 8, 1865 (2017).

29. Martin, L.-A. et al. An in vitro model showing adaptation to long-term oestrogen deprivation highlights the clinical potential for targeting kinase pathways in combination with aromatase inhibition. Steroids 76, 772–6 (2011).

30. Pisco, A. O. et al. Non-Darwinian dynamics in therapy-induced cancer drug resistance. Nat. Commun. 4, 2467 (2013).

31. Razavi, P. et al. The Genomic Landscape of Endocrine-Resistant Advanced Breast Cancers. Cancer Cell 34, 427–438.e6 (2018).

32. Wu, Y., Tamayo, P. & Zhang, K. Visualizing and interpreting single-cell gene expression datasets with Similarity Weighted Nonnegative Embedding. bioRxiv 276261 (2018). doi:10.1101/276261

33. Bausch-Fluck, D. et al. A mass spectrometric-derived cell surface protein atlas. PLoS One 10, e0121314 (2015).

34. Tang, D. G. Understanding cancer stem cell heterogeneity and plasticity. Cell Res. 22, 457–72 (2012).

35. Brooks, M. D., Burness, M. L. & Wicha, M. S. Therapeutic Implications of Cellular Heterogeneity and Plasticity in Breast Cancer. Cell Stem Cell 17, 260–71 (2015).

36. Chaffer, C. L. et al. Poised chromatin at the ZEB1 promoter enables breast cancer cell plasticity and enhances tumorigenicity. Cell 154, 61–74 (2013).

37. Chan, T. E., Stumpf, M. P. H. & Babtie, A. C. Gene Regulatory Network Inference from Single-Cell Data Using Multivariate Information Measures. Cell Syst. 5, 251–267.e3 (2017).

38. Liberzon, A. et al. The Molecular Signatures Database Hallmark Gene Set Collection. Cell Syst. 1, 417–425 (2015).

39. Aibar, S. et al. SCENIC: single-cell regulatory network inference and clustering. Nat. Methods 14, 1083–1086 (2017).

40. Grossman, R. L. et al. Toward a Shared Vision for Cancer Genomic Data. N. Engl. J. Med. 375, 1109–12 (2016).

41. Colleoni, M. et al. Annual Hazard Rates of Recurrence for Breast Cancer During 24 Years of Follow-Up: Results From the International Breast Cancer Study Group Trials I to V. J. Clin. Oncol. 34, 927–35 (2016).

42. Early Breast Cancer Trialists’ Collaborative Group (EBCTCG). Effects of chemotherapy and hormonal therapy for early breast cancer on recurrence and 15-year survival: an overview of the randomised trials. Lancet (London, England) 365, 1687–717 (2005).

43. Blows, F. M. et al. Subtyping of breast cancer by immunohistochemistry to investigate a relationship between subtype and short and long term survival: a collaborative analysis of data for 10,159 cases from 12 studies. PLoS Med. 7, e1000279 (2010).

44. Aceto, N. et al. Circulating Tumor Cell Clusters Are Oligoclonal Precursors of Breast Cancer Metastasis. Cell 158, 1110–1122 (2014).

45. Yu, M. et al. Circulating breast tumor cells exhibit dynamic changes in epithelial and mesenchymal composition. Science 339, 580–4 (2013).

46. Magnani, L. et al. SREBP1 drives KRT80-dependent cytoskeletal changes and invasive behavior in endocrine resistant ERα breast cancer. bioRxiv 380634 (2018). doi:10.1101/380634

47. Lawson, D. A. et al. Single-cell analysis reveals a stem-cell program in human metastatic breast cancer cells. Nature 526, 131–135 (2015).

48. Puram, S. V et al. Single-Cell Transcriptomic Analysis of Primary and Metastatic Tumor Ecosystems in Head and Neck Cancer. Cell 171, 1611–1624.e24 (2017).

49. Torre, E. et al. Rare Cell Detection by Single-Cell RNA Sequencing as Guided by Single-Molecule RNA FISH. Cell Syst. 6, 171–179.e5 (2018).

50. Butler, A., Hoffman, P., Smibert, P., Papalexi, E. & Satija, R. Integrating single-cell transcriptomic data across different conditions, technologies, and species. Nat. Biotechnol. 36, 411–420 (2018).

51. Lin, Y. et al. Evaluating stably expressed genes in single cells. bioRxiv 229815 (2018). doi:10.1101/229815

52. Lun, A. T. L., McCarthy, D. J. & Marioni, J. C. A step-by-step workflow for low-level analysis of single-cell RNA-seq data with Bioconductor. F1000Research 5, 2122 (2016).

53. Patel, A. P. et al. Single-cell RNA-seq highlights intratumoral heterogeneity in primary glioblastoma. Science 344, 1396–401 (2014).

54. Nguyen, Q. H. et al. Profiling human breast epithelial cells using single cell RNA sequencing identifies cell diversity. Nat. Commun. 9, 2028 (2018).

55. Durinck, S., Spellman, P. T., Birney, E. & Huber, W. Mapping identifiers for the integration of genomic datasets with the R/Bioconductor package biomaRt. Nat. Protoc. 4, 1184–91 (2009).

56. Langmead, B. & Salzberg, S. L. Fast gapped-read alignment with Bowtie 2. Nat. Methods 9, 357–9 (2012).

57. Li, H. et al. The Sequence Alignment/Map format and SAMtools. Bioinformatics 25, 2078–9 (2009).

58. Talevich, E., Shain, A. H., Botton, T. & Bastian, B. C. CNVkit: Genome-Wide Copy Number Detection and Visualization from Targeted DNA Sequencing. PLoS Comput. Biol. 12, e1004873 (2016).

59. Finak, G. et al. MAST: a flexible statistical framework for assessing transcriptional changes and characterizing heterogeneity in single-cell RNA sequencing data. Genome Biol. 16, 278 (2015).

60. Benjamini, Y., Hochberg, Y. & Benjaminit, Y. Controlling the False Discovery Rate: A Practical and Powerful Approach to Multiple Testing. Source: Journal of the Royal Statistical Society. Series B (Methodological) 57, (1995).

61. Sing, T., Sander, O., Beerenwinkel, N. & Lengauer, T. ROCR: visualizing classifier performance in R. Bioinformatics 21, 3940–1 (2005).

62. Breiman, L. Random forests. Mach. Learn. 45, 5–32 (2001).

63. Parker, J. S. et al. Supervised risk predictor of breast cancer based on intrinsic subtypes. J. Clin. Oncol. 27, 1160–7 (2009).

